# A helminth chitinase structurally similar to mammalian chitinase displays immunomodulatory properties

**DOI:** 10.1101/641837

**Authors:** Friederike Ebner, Katja Balster, Katharina Janek, Agathe Niewienda, Piotr H. Malecki, Manfred S. Weiss, Tara E. Sutherland, Arnd Heuser, Anja A. Kühl, Jürgen Zentek, Andreas Hofmann, Susanne Hartmann

## Abstract

Previously, we reported significant immunomodulatory effects of the entire excretory-secretory (ES) proteins of the first larval stage (L1) of the gastrointestinal nematode *Trichuris suis* in a rodent model of allergic hyperreactivity. In the present study, we aimed to identify the proteins accounting for the modulatory effects of the *T. suis* L1 ES proteins and thus studied selected components for their immunomodulatory efficacy in an OVA-induced allergic airway disease model. In particular, an enzymatically active *T. suis* chitinase mediated amelioration of airway hyperreactivity, primarily associated with suppression of eosinophil recruitment into the lung. The three-dimensional structure of the *T. suis* chitinase as determined by high-resolution X-ray crystallography revealed significant similarities to mouse acidic mammalian chitinase (AMCase). In addition, the unique ability of *T. suis* chitinase to form dimers, as well as acidic surface patches within the dimerization region may contribute to the formation of cross-reactive antibodies to the mouse homologs. This hypothesis is supported by the observation that *T. suis* chitinase treatment induced cross-reactive antibodies to mouse AMCase and chitinase-like protein BRP-39 in the AHR model. In conclusion, a biologically active *T. suis* chitinase exhibits immunomodulatory properties despite its structural similarity to the mammalian counterpart.

**Author summary:** Experimental immunotherapy via reintroduction of intestinal worms to treat and prevent autoimmune, chronic inflammatory or allergic diseases is being discussed but the underlying mechanisms are still not fully understood. Here, we investigated the immunomodulatory potential of specific proteins of the whipworm *Trichuris suis* that are secreted very early during larval development. Using a murine model of allergic lung disease, we show that in particular one *T. suis* protein, functionally characterized as an active chitinase, is reducing the lung inflammation. The *T. suis* chitinases three-dimensional protein structure revealed remarkable similarities to the hosts’ chitinase, an enzyme known to play a pivotal role in lung allergy. We also show that treatment with the helminth chitinase induced cross-reactive antibody responses against murine chitinase and chitinase-like proteins, both being inflammatory marker and regulators of type 2 immunity. Thus, our study provides a novel mechanism of immunomodulation by helminth components and may contribute to a better understanding of clinical responses of patients receiving helminthic therapy.

## Introduction

Chronic helminth infections are known to be accompanied by various immunomodulatory processes. The active modulation of host immune responses is believed to ensure long term persistence of the parasites and most likely involves proteins released by the worms (reviewed in (1–3)). Over the last decade, research has focused on identifying secreted helminth immunomodulators and mechanisms of suppression in order to understand natural infection and in parallel with the aim to exploit this knowledge for the benefit of unwanted human inflammatory diseases.

We and others have identified secreted products with immunosuppressive effects in autoimmune and inflammatory diseases from diverse parasitic helminths such as *Acanthocheilonema viteae* (4–11), *Schistosoma* (12–15), *Fasciola hepatica* (16,17), *Heligmosomoides polygyrus* (18,19), and *Wuchereria bancrofti* (20,21). Among the organisms currently explored for helminth-induced therapy of humans are the pig whipworm (*Trichuris suis*, TSO), the human hookworm (*Necator americanus*, NC), and also the human whipworm (*T. trichiura*, TTO). As *T. suis* can be easily maintained in pigs, but cannot multiply in humans, its administration as live parasite (TSO therapy) has been tested in a variety of clinical studies to improve human immune disorders (reviewed in (22)). Early success stories of treatments of ulcerative colitis (UC) and Crohn’s disease (CD) with TSO (23–25) were followed by larger trials with only modest results for various autoimmune diseases (22,26–28) and implicated the need for a greater understanding for the immunomodulatory mechanisms.

*T. suis* has a direct life cycle. Upon ingestion of larval stage 1 (L1) containing infective eggs, the larvae hatch and invade the mucosa of the large intestine and develop into adults by molting several times (29). When administered therapeutically to humans, there is evidence for a self-limiting, sterile colonization of *T. suis* (23), thus repeated TSO treatments are required. However, this transient exposure highlights the role of early larval stages to possess immunomodulatory properties. Using soluble products (*Ts*-SPs), or excretory-secretory proteins (*Ts*-ES) or isolated fractions from both adult worms and larval stages, several immunomodulatory mechanisms have been investigated (30–33). *Ts*-SPs from adult worms have been shown to interfere with activation of human dendritic cells and macrophages (31,34–37). We have reported on the immunomodulatory capacities of *Ts*-ES proteins from freshly hatched *T. suis* L1 *in vitro* by interfering with CD4^+^ T cell priming and *in vivo* by reducing clinical disease in a model of allergic airway disease (33). A recent study profiled *Ts*-ES proteins released by adult and larval stages of *T. suis*, and proved adult and larval (28-day larvae) *Ts*-ES possess anti-inflammatory properties. Indeed, the authors identified a subset of proteins with anti-inflammatory functions within adult *Ts*-ES (30). However, immunoregulators of the very early larval stages, the L1 of *T. suis* remain elusive.

Here, we concentrate on the identification and characterization of proteins selectively secreted by the first larval stages, the L1 of *T. suis*. We screened a set of recombinant larval proteins for immunomodulatory properties in a murine model of allergic airway disease and identified an enzymatically active *T. suis* chitinase that suppressed disease progression and recruitment of airway inflammatory cells. X-ray crystallography revealed the structure of the nematode enzyme to be highly similar to the host’s chitinases. Application of *T. suis* chitinase in the disease model induced cross-reactive antibodies to AMCase and chitinase-like protein BRP39. This implicated the contribution of *T. suis* chitinase, a structural similar enzyme to mammalian AMCase, in the immunomodulatory properties of secreted helminth proteins.

## Materials and Methods

### Animals

Female BALB/c mice were purchased from Janvier Labs (Le Genest-Saint-Isle, France). Mice were maintained at the Institute of Immunology, Department of Veterinary Medicine in Berlin and cared for in accordance with the principles outlined in the European Convention for the Protection of Vertebrate Animals used for Experimental and other Scientific Purposes and the German Animal Welfare Law. The mouse study was approved by the State Office of Health and Social Affairs Berlin, Germany (Landesamt für Gesundheit und Soziales; approval number G0144/10 and G0068/16).

### Murine OVA-induced allergic airway disease

Allergic airway inflammation was induced as previously described (4). Briefly, female BALB/c mice (8 weeks old) were sensitized on days 0 and 14 with 20 µg ovalbumin protein (OVA, grade VI, Sigma-Aldrich Chemie GmbH, Munich, Germany) emulsified in 2 mg alum (Imject™ Alum, Thermo Fisher Scientific, IL, USA). Non-allergic control animals (PBS) received equal volumes of PBS on days 0 and 14 instead of OVA. Mice were treated with 20 µg recombinant, intact *T. suis* proteins or heat inactivated proteins on days 0, 7 and 14 i.p., or left untreated. Intranasal OVA challenge (50 µg OVA/25 µl PBS) was performed on all groups of mice on days 28 and 29. Where indicated, lung function was assessed on day 30 at the Max Delbrück Center for Molecular Medicine (MDC) in Berlin. On day 31, mice were sacrificed, bronchoalveolar lavage (BAL) was performed and samples were taken for analysis.

### Airway hyperreactivity

Airway resistance and lung function was assessed via double chamber plethysmography (emka Technologies, Paris, France) in response to increasing doses of acetyl-ß-methylcholine (Metacholine, Sigma-Aldrich Chemie GmbH, Munich, Germany).

Mice were placed into a restrainer without anesthesia, adjusted (5 min) and the following protocol was applied: baseline measurement (4min), PBS (1 min), measurement (4 min), 6.25 mg/ml methacholine (1 min), measurement (4 min), 12.5 mg/ml methacholine (1 min), measurement (4 min), 25 mg/ml (1 min) methacholine, measurement (4min), 50 mg/ml methacholine (1 min), measurement (4 min).

Data were sampled once per second. For data analysis, averages of 5 sec intervals were used. Airway resistance (S_raw_) and expiratory time values were generated and analyzed using iox2 Software (iox v2.8.0.13, EMKA Technologies). Resulting data points were averaged again per mouse and methacholine dose. Data points with success rate < 50% and beat number <10 were excluded from the analysis.

### Bronchoalveolar lavage

Intratracheal bronchoalveolar lavage (BAL) was performed on i.p. anaesthetized mice by inflating the lungs with 800 µl PBS containing protease inhibitor cocktail (cOmplete™, Mini, EDTA-free Protease Inhibitor, Roche Diagnostics GmbH, Mannheim, Germany). The 1^st^ BAL was pelleted and the supernatant was stored at −20°C for cytokine/chemokine detection. Four additional lavages (4× 800 µl PBS) were performed for cell isolation and pelleted by centrifugation. All cells were pooled in 500 µl PBS and counted using a Neubauer cell chamber. 2× 100 µl cell suspension per sample were used for cytospin preparation. Object slides were dried and stained with Diff-Quick solutions (Labor und Technik Eberhard Lehmann GmbH, Berlin, Germany) for differential cell counts. Residual cells were used for flow cytometry.

### Histology and immunohistochemistry

Following lung lavage, the left lobe was split into two pieces. The lower tissue was degassed and fixed in a formalin solution (Roti-Histofix 10%, Carl Roth GmbH + Co. KG, Karlsruhe, Germany) for 6 h at room temperature and then stored at 4 °C. The tissue was embedded in paraffin and sections were stained with an antibody against RELM-α (polyclonal rabbit anti-mouse, Abcam, Cambridge, UK). Nuclei were stained with haematoxylin (Merck). Interstitial RELM-α^+^ cells were quantified in 5 HPF (high power fields, 400x magnification). Pictures are presented at 400x magnification. For evaluation of histomorphological changes, paraffin sections were stained with hematoxylin and eosin (H&E) and scored for inflammation and goblet cell hyperplasia. Briefly, score 1 for minor perivascular inflammation, score 2 for moderate perivascular and peribronchial inflammation and minimal goblet cell hyperplasia, score 3 for increased perivascular and peribronchial inflammation and increased goblet cell hyperplasia (beginning in smaller airways) and score 4 for severe perivascular, peribronchial and interstitial inflammation with goblet cell hyperplasia in smaller and larger airways.

### Multiplex assay

Bronchoalveolar lavage supernatants were analyzed for cytokine and chemokine levels using the Mouse Cytokine & Chemokine 26-plex ProcartaPlex™ Panel (ThermoFisher, MA, USA) according to the manufacturer’s instructions. The MAGPIX™ instrument (Luminex) was used and data were analyzed with Procarta Plex™ Analyst 1.0.

### Flow cytometry

The following monoclonal fluorochrome conjugated anti-mouse antibodies were used: CD11b (clone M1/70, APC-Cy7), CD64 (clone X54-5/7.1, PE-Cy7), IgG1 (clone RMG1-1, APC), IgG2a (clone RMG2a-62, PerCP-Cy5.5) were purchased from BioLegend, CA, USA. CD45 (clone 30-F11, eF450), MHC-II (clone M5/114.15.2, PE-Cy5), Gr1 (clone RB6-8C5, PE-Cy7), EpCAM (clone G8.8, APC) were purchased from eBioscience, CA, USA. CD11c (clone HL3, FITC), MHC-II (clone M5/114.15.2, PE), SiglecF (clone E50-2440, PE) were purchased from BD Biosciences, CA, USA. Viability dye in eFluor 506, eF660, eF780 and Streptavidin (FITC) were purchased from eBioscience, CA, USA. Polyclonal biotynilated anti-mouse Relm-α was purchased from Peprotech, NJ, USA. Samples were acquired using Canto II (BD Biosciences, CA, USA) and analyzed using Flow Jo software v9, Tristar.

### Quantitative RT-PCR

Following lung lavage, the upper right lobe was snap-frozen in liquid nitrogen and stored at −80 °C. RNA-Isolation from lung tissues was performed using the innuPREP RNA Mini Kit (Analytic Jena AG, Jena, Germany) according to manufacturer’s instructions. Reverse transcription of 1 µg RNA into cDNA was done using the High-Capacity cDNA Reverse Transcription Kit (Applied Biosystems, CA, USA). RT-PCR was performed on the LightCycler^®^ 480 instrument using the LightCycler^®^ 480 SYBR Green I Master mix (Roche Diagnostics, Mannheim, Germany) and oligonucleotides listed in Table 1. Gene expression is described relative to the housekeeping gene peptidylprolyl isomerase A (PPIA) and normalized to mRNA levels of the PBS group if not indicated differently.

**Table 1.**
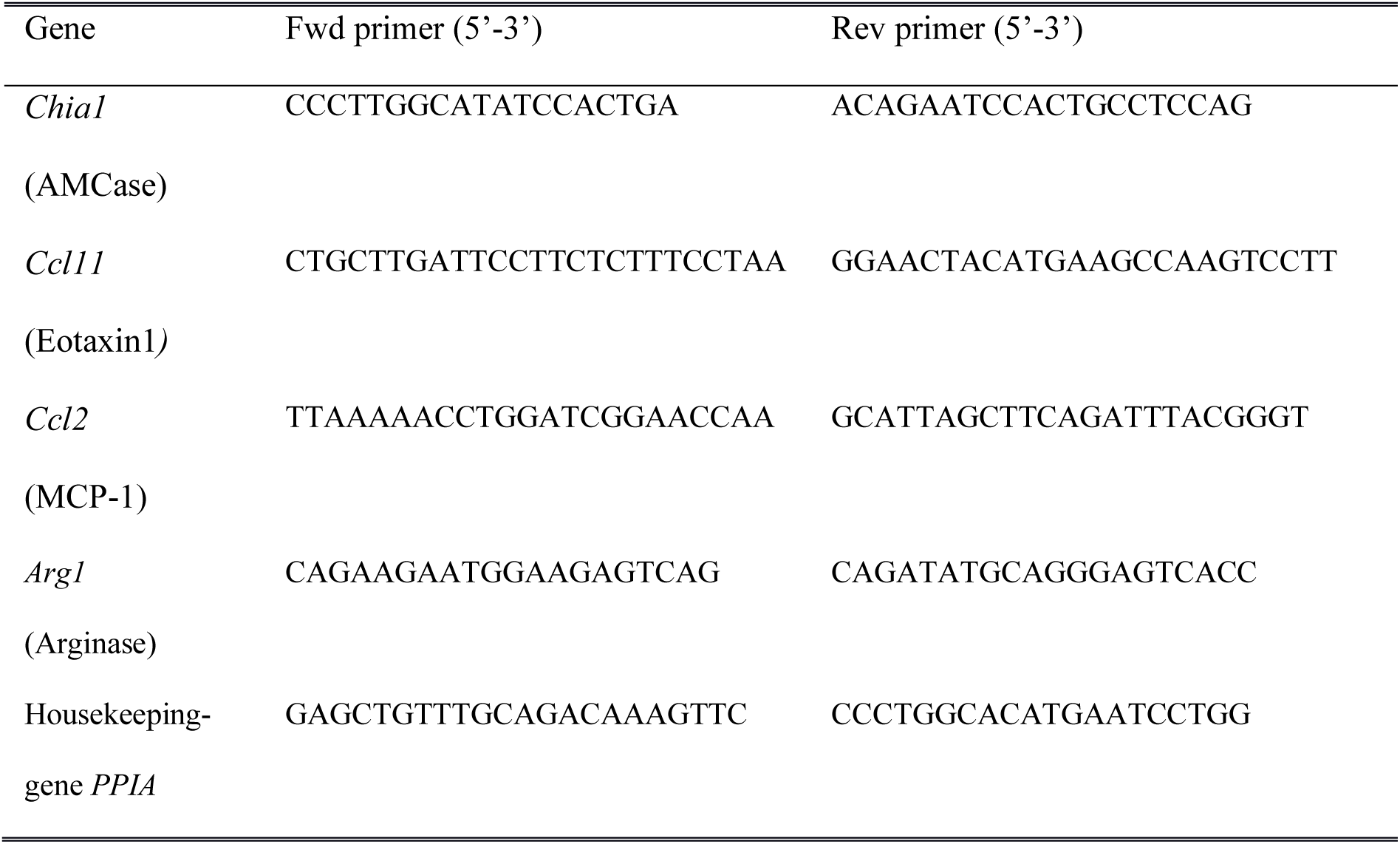
Sequences of oligonucleotides

### Isolation of *T. suis* larvae and production of excretory-secretory (ES) proteins

The *T. suis* life cycle was maintained in pigs (*Sus scrofa*) and the generation of larval material approved by the State Office of Health and Social Affairs Berlin, Germany (Landesamt für Gesundheit und Soziales; approval number H0296/12). Pigs were orally infected with 8000 embryonated *T. suis* eggs in 2 ml NaCl (0.9%). Following euthanasia on days 10, 18, 28 dpi *T. suis* larvae were manually collected from the caecum and colon. Larvae were washed 4 times for 15 min each in RPMI-1640 (PAN-Biotech, Aidenbach, Germany) supplemented with 200 U/ml penicillin, 200 µg/ml Streptomycin, 1.35 µg/ml Amphotericin B (all purchased from PAN-Biotech) by sedimentation and incubated overnight in RPMI-1640 supplemented with 1% Glucose (Sigma-Aldrich, St. Louis, US), 100 U/ml Penicillin, 100 µg/ml Streptomycin and 0.625 µg/ml Amphotericin B. Culture medium was replaced and larvae were incubated for another 5 days while conditioned media was collected every 2^nd^ day, sterilized by filtration (0.45 µm Minisart Syringe Filter, Sartorius AG, Göttingen, Germany) and stored at −20 °C. *T. suis* L1 ES proteins were isolated from freshly hatched larvae as previously described (33). Identification of *T. suis* larvae ES proteins (*in vitro* hatched L1, 10 dpi, 18 dpi, 28 dpi) was done by mass spectrometry as previously described (33). In brief, TCA-precipitated dried protein pellets of *T. suis* were suspended, reduced, alkylated and tryptic digested. The resulting peptides were analyzed by LC-MSMS on a 4700 proteomics Analyzer (ABSCIEX, Framingham, MS). Database searches were performed with Mascot software against NCBI database 20150331 (64057457 sequences). Proteins were accepted as identified if at least two MS/MS spectra provided a MASCOT score for identity (p<0.01).

### Recombinant protein expression

*T. suis* mRNA was isolated out of 10-20 mg worms by using the innuPREP RNA Mini Kit (Analytic Jena AG, Jena, Germany) according to manufacturer’s instructions. Reverse transcription of 1 µg RNA into cDNA was done using the Transcriptor First Strand cDNA Synthesis Kit (Roche Diagnostics GmbH, Mannheim, Germany).

The genes corresponding to selected *T. suis* proteins were amplified and cloned into the vector pLEXSYsat2 of the Leishmania Expression System (LEXSYcon2 Expression Kit, Jena Bioscience GmbH, Jena, Germany). The cloning vector was sequenced for verification. *Leishmania tarentolae* cells were transfected via electroporation (Nucleofector™ 2b Device, AmaxaTM Human T Cell Nucleofector™ Kit, Lonza, Basel, Switzerland) with the respective expression cassette containing the selective antibiotic marker Nourseothricin (NTC) and the *T. suis* protein coding gene. Transfected *L. tarentolae* cells were polyclonally selected and maintained as described in the manual.

### Protein purification

His_6_-tag fused, recombinant proteins secreted by transfected *Leishmania* cells were purified by liquid chromatography over His-Trap™ excel columns (GE Healthcare, Buckinghamshire, United Kingdom). Protein-containing fractions were pooled and dialysed two times against PBS (4 h and overnight). Finally, protein concentrations were assessed using Pierce™ BCA Protein Assay Kit (ThermoFisher Scientific, Rockford, USA). Identity of purified, recombinant proteins was verified by LC-MSMS analysis after tryptic digestion.

### SDS-PAGE and Western Blot

To validate recombinant expression of *T. suis* proteins, a 12% SDS gel was loaded with 1-2 µg purified, recombinant *T. suis* proteins and developed using silver staining (Pierce SilverStain kit, ThermoFisher Scientific, Rockford, US).

To visualize recombinant murine chitinases and CLPs, a 12% SDS gel was loaded with 20 µl cell culture supernatant of transfected COS-cells, 0.25 µg recombinant *T. suis* chitinase and 0.2 µg p53. Proteins were transferred to a nitrocellulose membrane (Amersham Protran 0.45 µm NC, GE Healthcare, Buckinghamshire, UK) using a semi-dry Western Blot system. Membranes were blocked for 1 h at room temperature in blocking buffer (5% BSA, PBS, 0.1% Tween). For detection of recombinant mouse chitinases, CLPs, p53 and *Ts*-Chit, an anti-His_6_-Tag antibody (1:200, Dianova GmbH, Hamburg, Germany) was used and, where indicated, pooled mouse serum (from *n* = 4 animals) at a 1:100 dilution in blocking buffer, followed by washing (PBS, 0.1% Tween) and incubation with goat anti-mouse IgG-HRP (1:2500) for 1 h at room temperature. After washing, blots were developed using WesternBright ECL (advansta, CA, USA) according to the manufacturer’s instructions and images were recorded with the chemiluminescence system Fusion SL 3500 WL (peqlab - VWR, Erlangen, Germany).

### Chitinase assay

Chitinase activity was analyzed by using the substrate and standard solutions of the fluorimetric chitinase assay kit (Sigma-Aldrich Chemie GmbH, Munich, Germany). Substrates 4-MU-(GlcNAc)_2_ (Exo-Chitinase activity) and 4-MU-(GlcNAc)_3_ (Endo-Chitinase activity) were adjusted to 1100 µM in DMSO and diluted 1:20 in assay buffer (100 mM citric acid, 200 mM sodium phosphate, pH 5.6) immediately prior to starting the assay by adding 50 µl of diluted substrate solution to 10 µl of protein KFD48490.1 (*Ts*_Chit) (5 µg/ml). The reaction mix was incubated for 30 min at 37 °C, and then quenched by adding 500 µl stop solution (500 mM sodium carbonate, 500 mM sodium bicarbonate, pH 10.7). 250 µl of the reaction solution were transferred in duplicates into a black 96-well plate and fluorescence of liberated 4-MU was measured using an excitation wavelength of 360 nm and recording emission at 450 nm. The effect of pH on KFD48490.1/*Ts*_Chit chitinase activity was determined by using assay buffers adjusted to pH = 2, 3, 4, 5, 6, 7, 8, 9 and 10. Heat stability of *Ts*_Chit chitinase was assessed by pre-incubation of the protein for 60 min at different temperatures (4°C – 90°C) followed by the chitinase assay.

### Crystallization and X-ray data collection

The recombinant protein solution was dialyzed after initial purification in 20mM HEPES buffer (pH 7.5) and purified over a HiLoad 16/600 Superdex 200 pg column with a HEPES running buffer (20 mM HEPES pH 7.7, 200 mM NaCl, 1 mM TCEP). Elution fractions were concentrated using Amicon Ultra-4 concentrator tubes (30,000 MWKO) to a final concentration of 33 mg/ml. A Gryphon pipetting robot was used to set up crystallization plates using the JSCG-plus Screen by mixing 0.25 µl protein solution with 0.25 µl crystallization cocktail solution. Plates were sealed and incubated at 20°C. Crystal growth was monitored regularly under an optical microscope. The crystals appeared in condition H7 (0.8 M ammonium sulfate, 0.1 M BIS-Tris pH 5.5, 25% (w/v) PEG 3350). Protein crystals were harvested, cryo-protected using reservoir solution supplemented with 25% (v/v) ethylene glycol and stored in liquid nitrogen. Diffraction data were collected on BL14.1 at the BESSY II synchrotron of the Helmholtz-Zentrum Berlin (38) and processed using XDSAPP (39). Relevant processing statistics are shown in Table 2. The structure was solved by molecular replacement using the structure of Human Chitotriosidase (CHIT1) as a search model (PDB-Id 5HBF, (40)). After several cycles of manual rebuilding using COOT (41) and refinement using REFMAC5 (42), refinement converged at R-factors of 19 and 23 % for the working R and the free R, respectively (Table 3). The refined structure features good refinement statistics and excellent geometric parameters. The refined coordinates and the associated structure factor amplitudes were deposited in the PDB using the accession code 6G9C.

**Table 2.** MALDI-TOF MS verified T. suis L1 ES proteins that were selected and recombinantly expressed using LEXSY

| NCBI-ID | MW<br>(kDa) | Known homology | N-Glycosylation | O-Glycosylation |
| --- | --- | --- | --- | --- |
| KFD48490.1 | 55.45 | Acidic mammalian chitinase<br>[*T. trichiura*] | - | 28 |
| KFD55729.1 | 13.89 | - | - | 1 |
| KFD54802.1 | 31.06 | CAP domain containing protein<br>[*T. trichiura*] | 3 | 10 |
| KFD45500.1 | 37.04 | venom allergen 5<br>[*T. trichiura*] | 1 | 24 |
| KFD47618.1 | 14.18 | hypothetical protein<br>[*T. trichiura*] | - | - |
| KFD53667.1* | 31.06 | hypothetical protein<br>[*T. trichiura*] | 5 | - |

Sequence similarity search for protein homology was performed using BLAST/NCBI. Glycosylation prediction was performed using NetOGlyc 4.0.0.13 for O-Glycosylation and NetNGlyc 1.0 Server for N-Glycosylation. \* formerly identified using HelmDB database and recombinantly expressed based on this sequence.

**Table 3.**
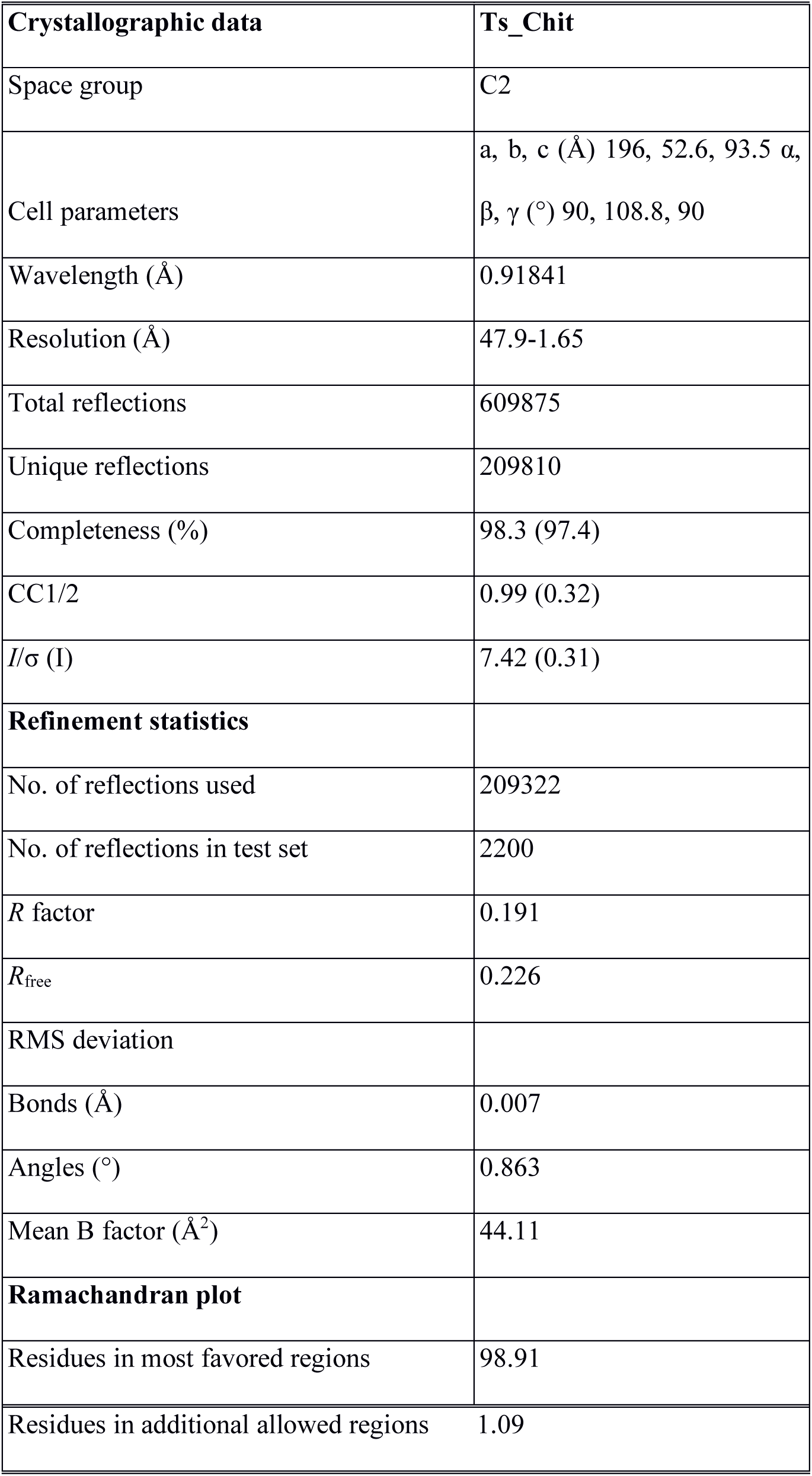
X-ray-based structure analysis data collection and refinement statistics

| Crystallographic data | Ts_Chit |
| --- | --- |
| Space group | C2 |
| Cell parameters | a, b, c (Å) 196, 52.6, 93.5 $\alpha$,<br>$\beta$, $\gamma$ (°) 90, 108.8, 90 |
| Wavelength (Å) | 0.91841 |
| Resolution (Å) | 47.9-1.65 |
| Total reflections | 609875 |
| Unique reflections | 209810 |
| Completeness (%) | 98.3 (97.4) |
| CC1/2 | 0.99 (0.32) |
| $I/\sigma(I)$ | 7.42 (0.31) |
| **Refinement statistics** |  |
| No. of reflections used | 209322 |
| No. of reflections in test set | 2200 |
| $R$ factor | 0.191 |
| $R_{\text{free}}$ | 0.226 |
| RMS deviation |  |
| Bonds (Å) | 0.007 |
| Angles (°) | 0.863 |
| Mean B factor (Å<sup>2</sup>) | 44.11 |
| **Ramachandran plot** |  |
| Residues in most favored regions | 98.91 |
| Residues in additional allowed regions | 1.09 |

### Structure based sequence alignment and surface electrostatics

To obtain an overview of structural similarities and differences between *Ts*-Chit and host chitinases, the topology and secondary structure elements of *Ts*-Chit were compared to six mouse chitinases and chitinase-like proteins by means of a structure-based amino acid sequence alignment (see Supplementary Information S2). Secondary structure elements were predicted with PSIPRED (43) and a structure-based amino acid sequence alignment was constructed using SBAL (44).

In order to assess whether *Ts*-Chit possesses any features that might serve as a potential molecular mimicry of host chitinases when acting as an antigen, we undertook a comparison of surface features of *Ts*-Chit with mouse chitinases. In general, this approach is limited by the availability of only one experimental 3D structure of mouse chitinases. Using the amino acid sequences of three mouse chitinases (*M. musculus* AMCase, Chit1 isoform 1 and Ym1), known 3D structures or close structural homologues of these proteins were identified in the PDB using pGenThreader (45), revealing mouse AMCase as the only mouse protein suitable for comparison in this context.

Using the monomer and dimer structures of *Ts*-Chit as well as the structure of mouse AMCase (PDB 3FY1), surface electrostatics for these three proteins were calculated with APBS (46), and visualized with UCSF Chimera (47).

### Statistical analysis

Statistical analysis was done using GraphPad Prism software v7 (GraphPad Software, Ca, USA) using Kruskal-Wallis test (Dunn’s multiple comparisons test), Mann-Whitney test or unpaired Student’s *t*-test as indicated. Values with *p* < 0.05 (*), *p* < 0.005 (**), *p* < 0.001 (***) were considered to be significant.

## Results

### *T. suis* L1 stage-specific excretory-secretory protein KFD48490.1 affects allergic airway inflammation

We previously showed that *T. suis* excretory/secretory proteins (ES proteins) collected from L1 hatched *in vitro* reduced clinical signs of murine allergic airway disease (33). To identify the early released immunomodulatory proteins responsible for this effect, we now investigated the protein composition of L1 ES proteins and compared it to L2, L3 and L4 larval stages of *T. suis* released products. For stage-specific expression patterns, parasites were isolated from pigs infected with 8000 TSO at 10, 18 and 28 days post-infection (resulting in L2, L3, and L4 larvae, respectively) and cultivated to collect conditioned media for mass spectrometry. Alternatively, eggs were hatched *in vitro* to generate L1 larvae and conditioned media, as previously described (33). Comparative proteomic analysis of *T. suis* ES proteins **(Fig. 1A and Supplementary Table S1)** revealed some overlap between first stage (*in vitro* hatched) L1 larvae and L2 larvae (10 day larvae), but no overlap of L1 ES proteins with later stages of larval development (L3 ES - 18 day larvae and L4 ES - 28 day larvae). Four L1-specific proteins and two L1/L2-specific proteins with predicted signal sequences and a lack of transmembrane domains (**Table 2**) were recombinantly expressed. Since glycosylation can be crucial for the immunomodulatory effects of secreted helminth proteins, we used the eukaryotic expression system LEXSY, based on the protozoan *L. tarentolae* (**Fig. 1B**). Notably, selected *T. suis* L1 proteins were distinct in size (ranging from 13 to 56 kDa, **Fig. 1C**), predicted glycosylation sites and homology-based annotation (**Table 2**).

**Figure 1.**
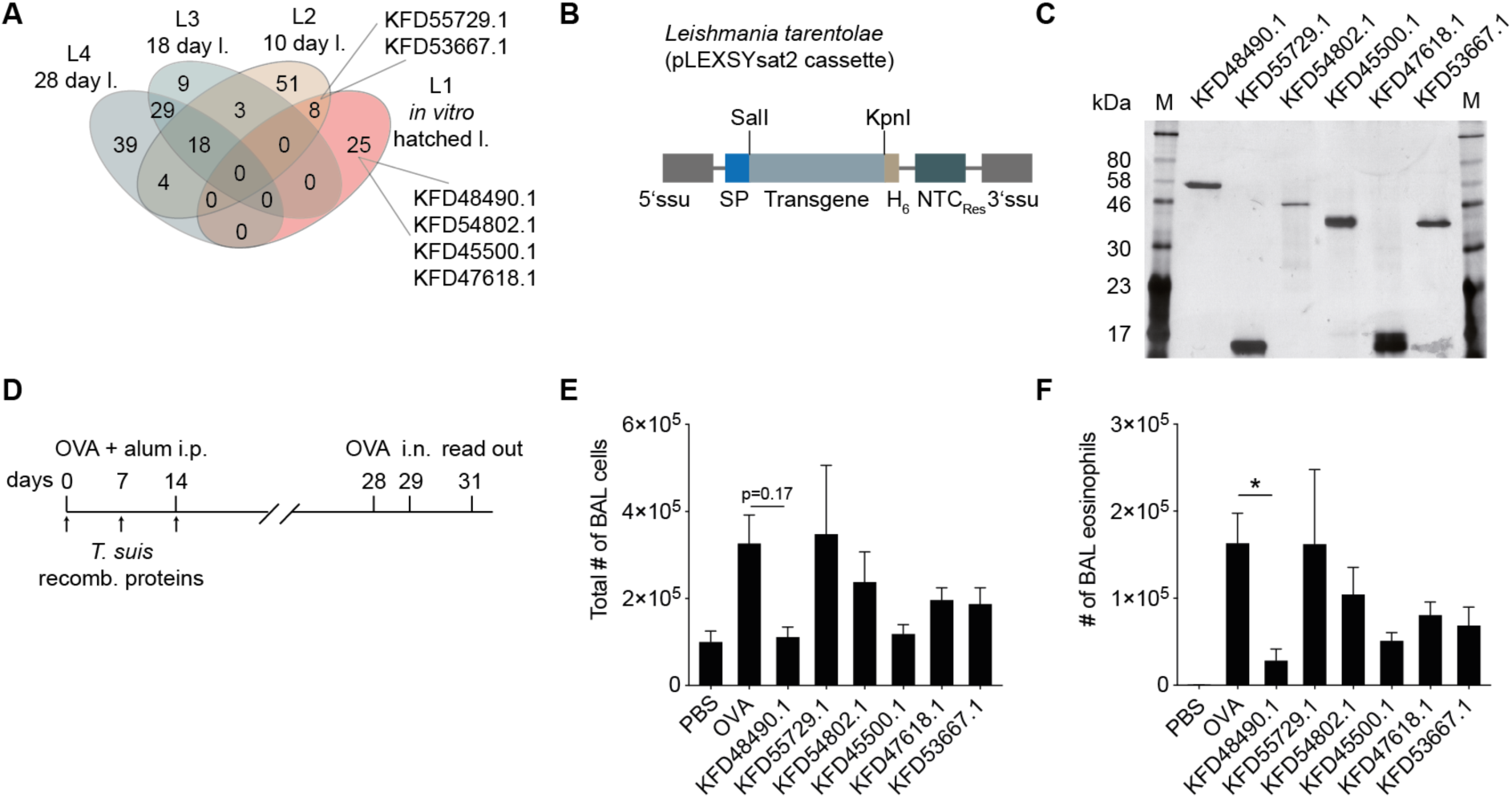
*T. suis* L1 ES proteins are highly stage-specific and candidates reduce allergic airway hyperreactivity when recombinantly expressed and applied. (**A**) Protein ID counts resulting from proteomic MS-based analysis (see **Supplementary Table S1**) were compared across the proteomic datasets of *T. suis* L1, L2, L3 and L4 ES proteins. Shown are protein IDs selected for recombinant expression. (**B**) Expression cassette of LEXSY eukaryotic expression system illustrating positioning of the export signal peptide that mediates secretion (SP), the selection antibiotic (nourseothricin, NTC) and the His_6_-tag for purification (H_6_). The cassette was integrated into the *L. tarentolae* genome. (**C**) Silver-stained SDS/PAGE gel of recombinantly expressed *T. suis* L1 proteins. (**D**) Experimental set-up of acute ovalbumin-(OVA)-induced allergic airway hyperreactivity model and treatment regimen with recombinant *T. suis* proteins. Animals were OVA-sensitized on days 0 and 14 and challenged with OVA intranasally on days 28 and 29. BAL fluid was collected from pretreated or untreated animals at day 31. (**E**) Total number of cellular infiltrates in BAL fluid. Shown are non-allergic mice (PBS controls), untreated allergic mice (OVA) and treatments with recombinant *T. suis* proteins indicated by GenBank IDs. (**F**) Numbers of eosinophils in BAL fluid detected by differential cell counts of cytospin preparations stained with DiffQuick over the different treatment groups. Data shown are mean + SEM of *n* = 5–10 animals from 2 separate experiments. Statistically significant differences were determined using a Kruskal-Wallis test (Dunn’s multiple comparisons test) and are indicated, * *p* ≤ 0.05.

We initially screened for immunomodulatory effects of the six recombinant *T. suis* L1 proteins using the OVA (ovalbumin)-induced allergic airway hyperreactivity model in female BALB/c mice. Mice were treated with recombinant *T. suis* proteins during sensitization with the allergen OVA on days 0, 7 and 14 (**Fig. 1D**). The numbers of total BAL infiltrates and BAL eosinophils were detected 2 days after the 2^nd^ intranasal OVA challenge at day 31 (**Fig. 1D**) and analyzed in parallel to control mice lacking OVA sensitization, but receiving OVA-challenge (PBS) and untreated allergic mice (OVA). Analysis of total cell numbers and cellular composition of the BAL fluid found that KFD48490.1 (homologous to *T. trichiura* acidic mammalian chitinase) and KFD45500.1 (homologous to *T. trichiura* venom allergen 5) reduced the total numbers of cellular BAL fluid infiltrates, but only KFD48490.1 treated mice showed a significant reduction of eosinophils infiltrating the lungs in comparison to untreated allergic (OVA) mice (**Fig. 1 E and F**).

### Immunoregulatory KFD48490.1 is an active chitinase of *T. suis* L1 larvae

Based on amino acid sequence similarity, database research predicted KFD48490.1 was homologous to acidic mammalian chitinase (AMCase) of *T. trichiura* (**Table 2**). The amino acid sequence of KFD48490.1 revealed the presence of the catalytic motive DxDxE (*T. suis* DLDWE, **Fig. 2A**), typically observed for active chitinases, and a C-terminal chitin-binding domain with cysteine residues responsible for the chitin binding (**Fig. 2A**). Alignment studies additionally proved high sequence similarities to other known helminth chitinases including a putative endochitinase from the pig infecting *Ascaris suum* and a glycosyl hydrolase from the human hookworm *N. americanus* (**Fig. 2B**). Active chitinases cleave chitin from the end of a 18 polymer chain (exochitinase) and/or hydrolyse small oligomers to generate N-acetylglucosamine monomers within the oligomer chain (endochitinase) (48). To examine whether KFD48490.1 exerts true chitinase activity, we used two different substrates, 4-MU-(GlcNAc)_3_ and 4-MU-(GlcNAc)_2_, allowing us to evaluate both endo- and exochitinase activity, respectively (**Fig. 2C**). KFD48490.1 clearly presented both, endo- and exochitinase activity, whereas extensive, heat-induced proteolysis resulted in a loss of chitinase activity (**Fig. 2C**).

**Figure 2.**
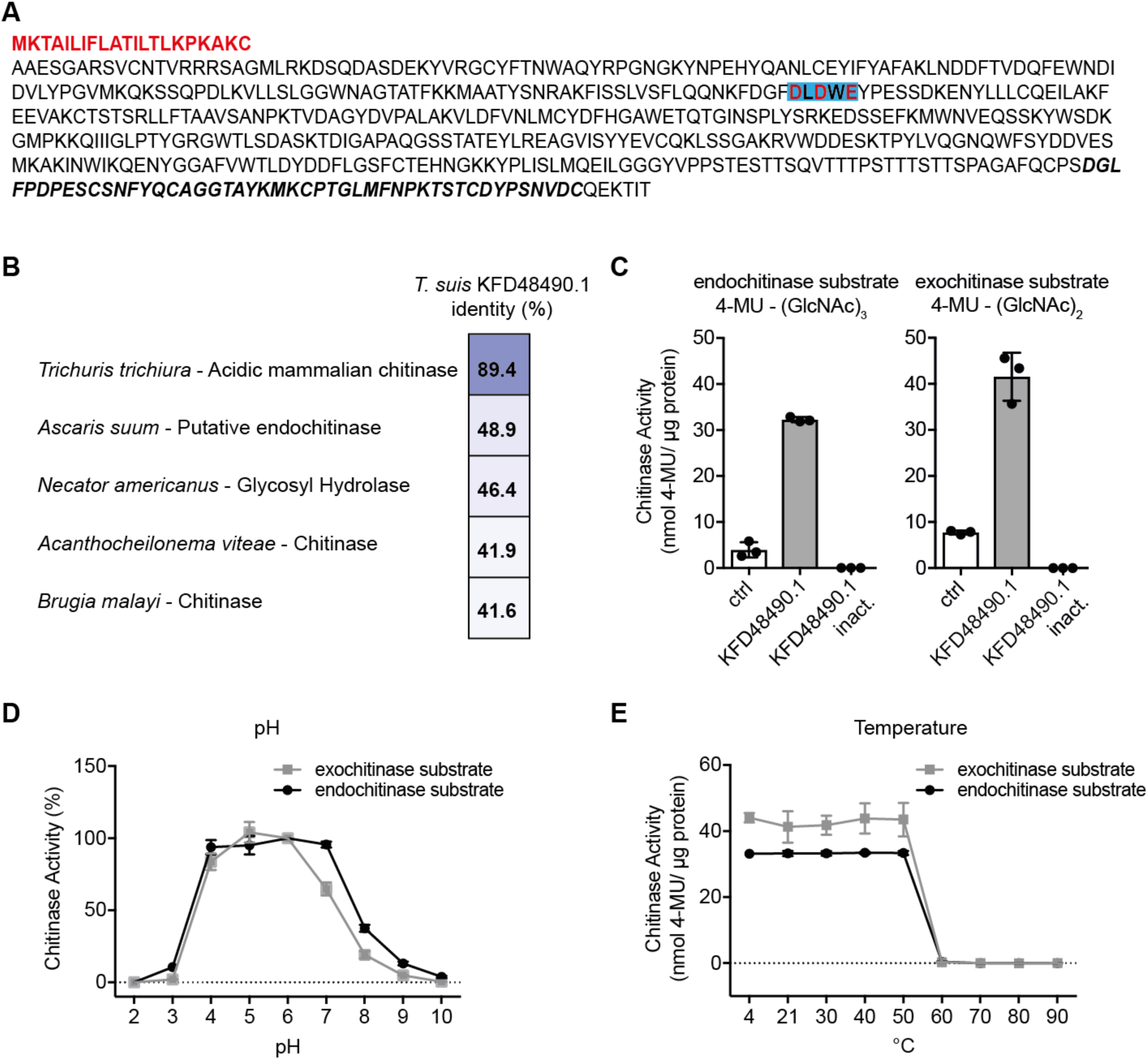
KFD48490.1 is an active chitinase. (**A**) KFD48490.1 amino acid sequence with mapped signal peptide (bold, red), catalytically active residues (highlighted in blue/bold red) and the chitin-binding domain (bold, black, italic). (**B**) Amino acid sequence alignment of KFD48490.1 and known nematode chitinases. (**C**) Fluorimetric chitinase assay testing recombinant KFD48490.1, heat-inactivated KFD48490.1 and a chitinase control enzyme (ctrl) with substrates for endochitinase (left) and exochitinase (right) activity. Data shown are mean ± SD of *n* = 3 replicates from one of three independent experiments. (**D**) Chitinase assay testing KFD48490.1 activity depending on pH (range 2-10). (**E**) Chitinase assay testing KFD48490.1 pre-incubated at different temperatures (4–90 °C) for 1 h prior to the assay. Data shown are mean ± SD of *n* = 3 replicates.

With regard to the biology of nematodes, we explored the influence of low pH and high temperatures hypothesizing that *T. suis* chitinase will be active at low pH and elevated temperatures in the pig’s intestine to support larval hatching and development (49,50). Interestingly, KFD48490.1 remained enzymatically active over a wide range of pH 4–7 (**Fig. 2D**) and up to temperatures of 50°C (**Fig. 2E**). Together, these data confirmed the functional annotation of KFD48490.1 as *T. suis* L1 true chitinase, and thus the protein is referred to as *Ts*-Chit from hereon.

### *T. suis* chitinase (*Ts*-Chit)-mediated regulation of airway hyperreactivity only partially depends on protein integrity

To describe the effects of *Ts*-Chit treatment on allergic airway disease we analyzed airway resistance, expiration time and BAL fluid profiles in more detail. In addition, we included a group of mice treated with heat-inactivated, recombinant *Ts*-Chit to assess whether immunomodulatory effects were linked to the enzymatically active *T. suis* chitinase protein. Treatment of animals with intact and active *Ts*-Chit significantly reduced the number of total cells (**Fig. 3A**) and eosinophils (**Fig. 3B**) present in BAL fluid 2 days after 2^nd^ OVA-challenge. Differential cell counts of BAL fluid further revealed that neutrophil numbers were largely unaffected (**Fig. 3C**). A trend towards an increase of alveolar macrophages cell numbers was observed in *Ts*-Chit compared to allergic (OVA) mice (**Fig. 3D**). Flow cytometry of BAL cells confirmed the reduced frequency of eosinophils (CD45^+^CD11b^+^GR1^low^CD11c^−^SiglecF^+^) in *Ts*-Chit treated compared to untreated (OVA) allergic animals (**Fig. 3E and F**), and also supported increased frequencies of alveolar macrophages, identified as CD11b^low^CD11c^+^ cells among CD45^+^ cells (**Fig. 3G**), as observed as a trend in differential cell counts. The respiratory responses resulting from *Ts*-Chit treatment were measured using non-invasive whole-body plethysmography 24 h after OVA challenge. Importantly, also the non-sensitized control group (PBS) was intranasally challenged with OVA. Preventative administration of *Ts*-Chit caused restoration of expiration time to the level of PBS control mice at baseline (**Fig. 3H**). A trend in recovery was also noticed for treating with heat-inactivated *Ts*-Chit (**Fig. 3H**). Measuring airway resistance to increasing doses of methacholine (MCh) revealed that *Ts*-Chit treatment attenuated exacerbation of MCh-induced airway hyperreactivity), but treatment with inactivated *Ts*-Chit also had some effects (**Fig. 3H, I and J)**. We next evaluated, whether the reduced cellularity and improved lung function observed in *Ts*-Chit treated animals was reflected in BAL cytokine levels. OVA sensitization and challenge caused a significant increase of the cytokines IL-4, IL-5 and IL-13 detected in BAL fluid; however, *Ts*-Chit treatment did not reduce but rather enhanced local Th2 cytokine production (**Fig. 3K**). Interestingly, *Ts*-Chit and heat-inactivated *Ts*-Chit alike accelerated IL-18 levels in BAL fluid (**Fig. 3K**), a cytokine known to induce IL-4, IL-5 and IL-13 release by mast cells and basophils (51). Thus, cellular infiltration and the respiratory response were improved in mice treated with *Ts*-Chit, but the Th2 cytokine response was unchanged or elevated in comparison to untreated allergic mice (OVA).

**Figure 3.**
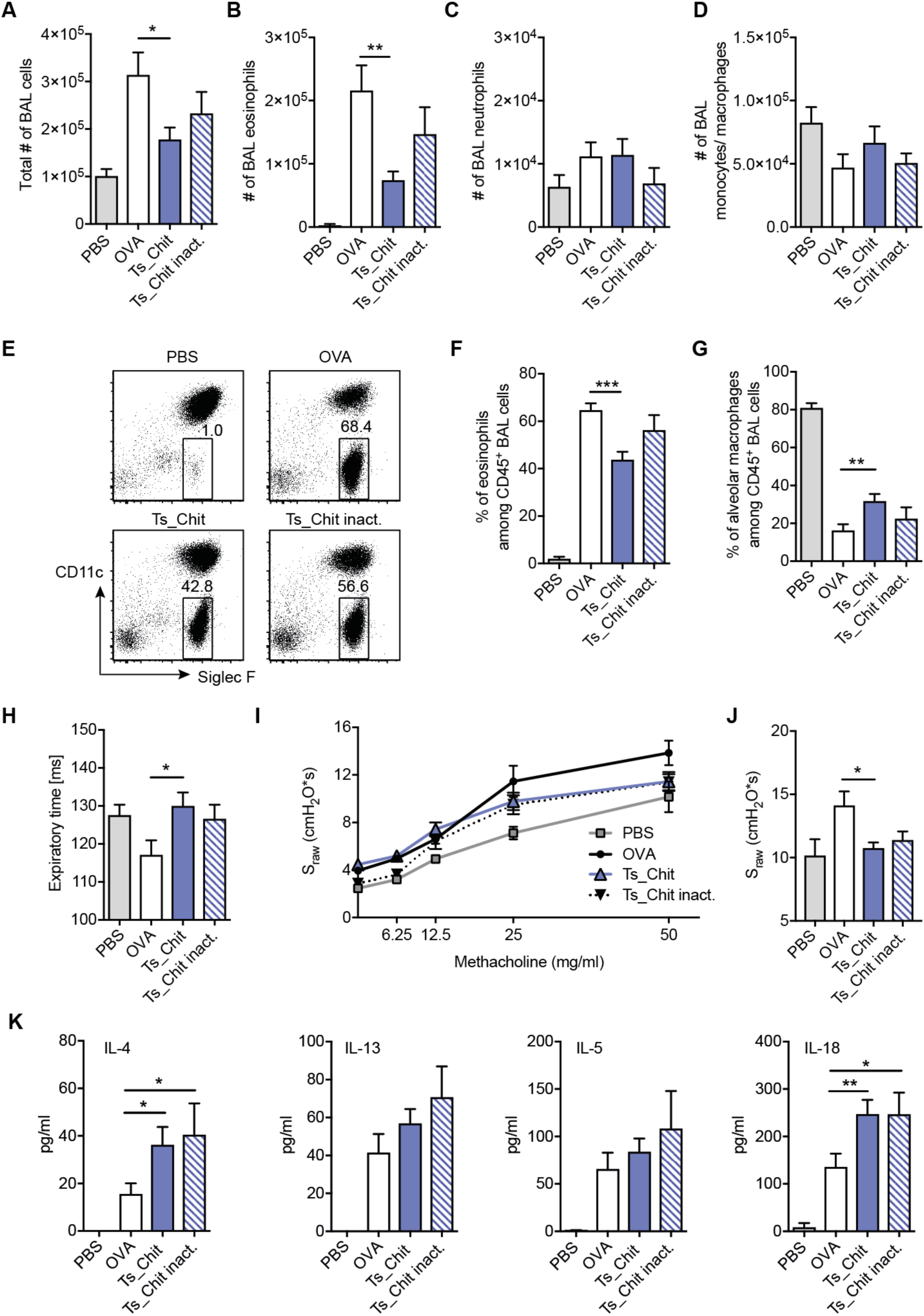
*Ts*_Chit treatment mediates attenuation of airway hyperreactivity. Bronchoalveolar lavage (BAL) was performed two days after OVA challenge and the BAL fluid was examined for (**A**) total number of infiltrating cells (Neubauer chamber), (**B**) BAL eosinophils, (**C**) BAL neutrophils and (**D**) monocytes/macrophages using cytospin preparations and DiffQuick staining. (**E**) Representative plots of BAL infiltrating cells using flow cytometry and focusing on eosinophil frequencies (identified as CD45^+^CD11b^+^GR1^low^CD11c^−^SiglecF^+^). Summary of flow cytometric analysis of (**F**) BAL eosinophils and (**G**) alveolar macrophages (CD45^+^CD11b^low^CD11c^+^). (**H**) Basal expiration time during PBS inhalation measured using whole-body double-chamber plethysmography 24 h after OVA challenge. (**I**) Concentration response curve of methacholine sensitivity. Specific airway resistance (S_raw_) was assessed in response to increasing doses of methacholine. (**J**) S_raw_ values recorded when nebulizing 50 mg/ml of methacholine. (**K**) BAL cytokines detected two days after challenge using bead-based multiplexing. **A-K** data are presented as mean ± SEM of *n* = 10–12 animals from *n* = 3 separate experiments. Statistical significance was assessed using a two-tailed Mann-Whitney test and results are indicated by * *p* ≤ 0.05, ** *p* ≤ 0.01, *** *p* ≤ 0.001.

To further delineate the direct effects of *Ts*-Chit treatment on lung inflammation of OVA-sensitized and challenged mice, we investigated lung tissue pathology and recruitment of alternatively activated macrophages (AAM). While histopathological findings remained overall similar in the lungs of untreated (OVA) and *Ts*-Chit treated allergic animals (**Fig. 4A**), immunohistochemistry revealed increased numbers of interstitial RELMα ^+^ cells, a signature marker for AAM that mediates lung vascularization and tissue repair (52,53) when mice were treated with *Ts*-Chit compared to OVA (**Fig. 4B and C**). A similar trend was observed for inactivated *Ts*-Chit treated animals. Since this was in contrast to our previous observation on *Ts*-ES L1-induced suppression of RELMα^+^ cells (33) and given that the resistin-like molecule (RELM) α is not only expressed in macrophages but also epithelial cells and tissue infiltrating eosinophils, we used smaller groups of mice to define Relmα expression to a interstitial macrophage phenotype by flow cytometry. Lung tissue macrophages were analyzed in homogenates of lavaged lungs and defined as EpCAM^−^CD11c^+^CD64^+^MHCII^+^SiglecF^−^ (**Fig. 4D**). While overall frequencies of tissue macrophages were not affected by *Ts*-Chit treatment (**Fig. 4E**), we again detected an increase in Relmα^+^ macrophages, but did not reach statistical significance (**Fig. 4F**). Contrary to the increase of Relmα^+^ cells, RT-PCR of homogenized lung tissue of all groups of mice revealed downregulation of *arginase-1* and *MCP-1* mRNA when mice were treated with intact and also inactivated *Ts*-Chit compared to PBS (**Fig. 4G**). Supporting reduced numbers of eosinophil infiltration upon *Ts*-Chit treatment, we detected downregulation of CCL11/eotaxin mRNA, a potent eosinophil chemoattractant that stimulates recruitment of eosinophils from the blood to sites of allergic inflammation (**Fig. 4E**). Together, these data illustrate that protein integrity of the *Ts*-Chit is only partially responsible for the treatment effects leading to attenuated airway hyperreactivity.

**Figure 4.**
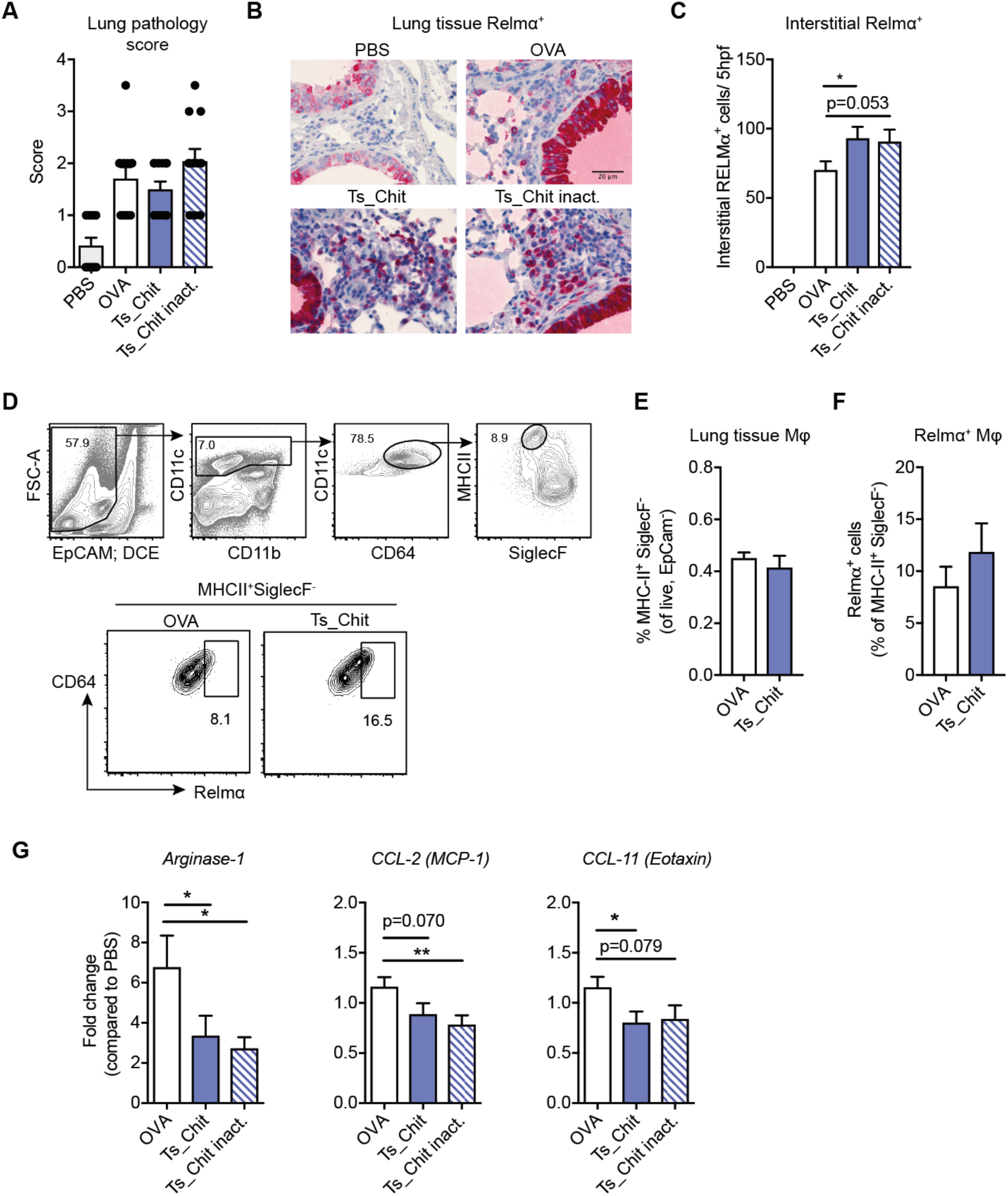
*Ts*-Chit-mediated effects on lung parenchymal macrophages. (**A**) Histopathological evaluation of tissue inflammation. (**B**) Histological sections of paraffin-embedded lung tissues from treated (*Ts*-Chit and *Ts*-Chit inact.), untreated (OVA) and non-sensitized (PBS) controls stained for RELMα (red) and haematoxylin (steelblue) and presented at 400× magnification. (**C**) Histological quantification of interstitial RELMα^+^ cells. (**D**) Gating strategy and representative plots of RELMα^+^ macrophages (EpCAM^−^CD11c^+^CD64^+^MHCII^+^Siglec^−^) of lung homogenates from OVA-sensitized and challenged mice with or without *Ts*-Chit treatment summarized in (**E**) and (**F**). (**G**) mRNA expression of *arginase-1, monocyte chemotactic protein 1 (MCP1/CCL-2)* and *CCL-11/Eotaxin* in lung tissue samples of mice from challenge experiments after lung lavage. Expression was normalized by using the housekeeper gene *peptidyl prolylisomerase A* (*PPIA*) and is presented as fold difference to the PBS control group. **A, C, G** data are presented as mean + SEM of *n* = 10–12 animals from *n* = 3 separate experiments. **D-F** data are generated from *n* = 4–5 mice from one experiment. Statistical significance was assessed using a two-tailed Mann-Whitney test and is indicated by * *p* ≤ 0.05, ** *p* ≤ 0.01.

### *T. suis* chitinase reveals high structural similarity to murine chitinase but presents dimeric structure

Given the results of the *in vivo* AHR experiments with heat-inactivated *Ts*-Chit, a direct immunomodulatory effect of *Ts*-Chit or a role for its chitinase activity was very unlikely. Direct immunomodulatory effects of ES or soluble proteins from adult and larval parasitic nematodes have previously been shown *in vitro* and various modulatory activities were found such as targeting pro-inflammatory cytokine release from bone-marrow derived dendritic cells or macrophages and induction of arginase-1, IL-10 and NO in macrophages (4,30,35,54,55), the suppression of antigen-specific or unspecific CD4^+^ T cell proliferation (56) or induction of Foxp3^+^ regulatory T cells (19,57). In contrast, recombinant *T. suis* chitinase failed to show strong immunomodulation in any of the above mentioned assays (data not shown).

We therefore examined the implications of *Ts*-Chit treatment on the family of murine chitinases and chitinase-like proteins (CLP), that are highly implicated in pathology of Th2-mediated allergic airway inflammation (58). Interestingly, *T. suis* chitinase was shown to have a high percentage of amino acid sequence identity when compared to the two true murine chitinases (AMCase (41.6%) and Chit1(41.5%)) or the murine CLPs Ym1 (39%), Ym2 (37.5%) or BRP-39 (36.7%) (Suppl. Fig. 2).

To better understand structural differences between nematode and mammalian chitinases and the structural basis of the serological response to *Ts*-Chit treatment, we determined the crystal structure of the nematode chitinase protein by X-ray crystallography. The final model analysis data collection and refinement statistics are summarized in **Table 3**. They indicate that the structure is of high resolution, that it is well refined to very good statistics and hence that it is of high quality. The overall structure reveals a TIM-barrel or (ßα)8-barrel motif, which means that 8 ß-strands form the central barrel which is surrounded by 8 α-helices (**Fig. 5A and C**). The active site with the sequence motif D_149_xD_151_xE_153_ is depicted in **Figure 5A.2**. The crystal structure furthermore revealed a P2-symmetric dimer of the protein, which is stabilized by an intermolecular disulfide bond formed by a cysteine residue at position 180 (**Fig. 5A.1 and B**). Such dimerization has not been described so far for human or murine chitinases, nor has the occurrence of a cysteine residue at position 180. In addition, the structural alignment with human AMCase (3FXY), human chitotriosidase (1GUV), human YKL-40 (1NWR) and murine Ym1 (1VF8) (**Fig. 5C**) indicates highly similar structural features in the core regions (marked in red) and certain areas, mostly at the protein surface, where the structure differs between nematode and mammalian chitinases (highlighted in green/blue). A detailed structure based sequence alignment of *T. suis* chitinase, mouse chitinases and CLPs is given in **Supplementary S2**. Notably, the C-terminal region of *Ts*-Chit from amino acid 401 to 495 contains many threonine repeats and is referred to as unordered region. The *T. suis* chitinase dimerization was further examined by comparing reducing (+ ß-mercaptoethanol) and non-reducing (− ß-mercaptoethanol) conditions for SDS/PAGE loading. We indeed detected a second, higher band at around 75 kDa indicative for dimerization under non-reducing conditions (**Fig. 5D**).

**Figure 5.**
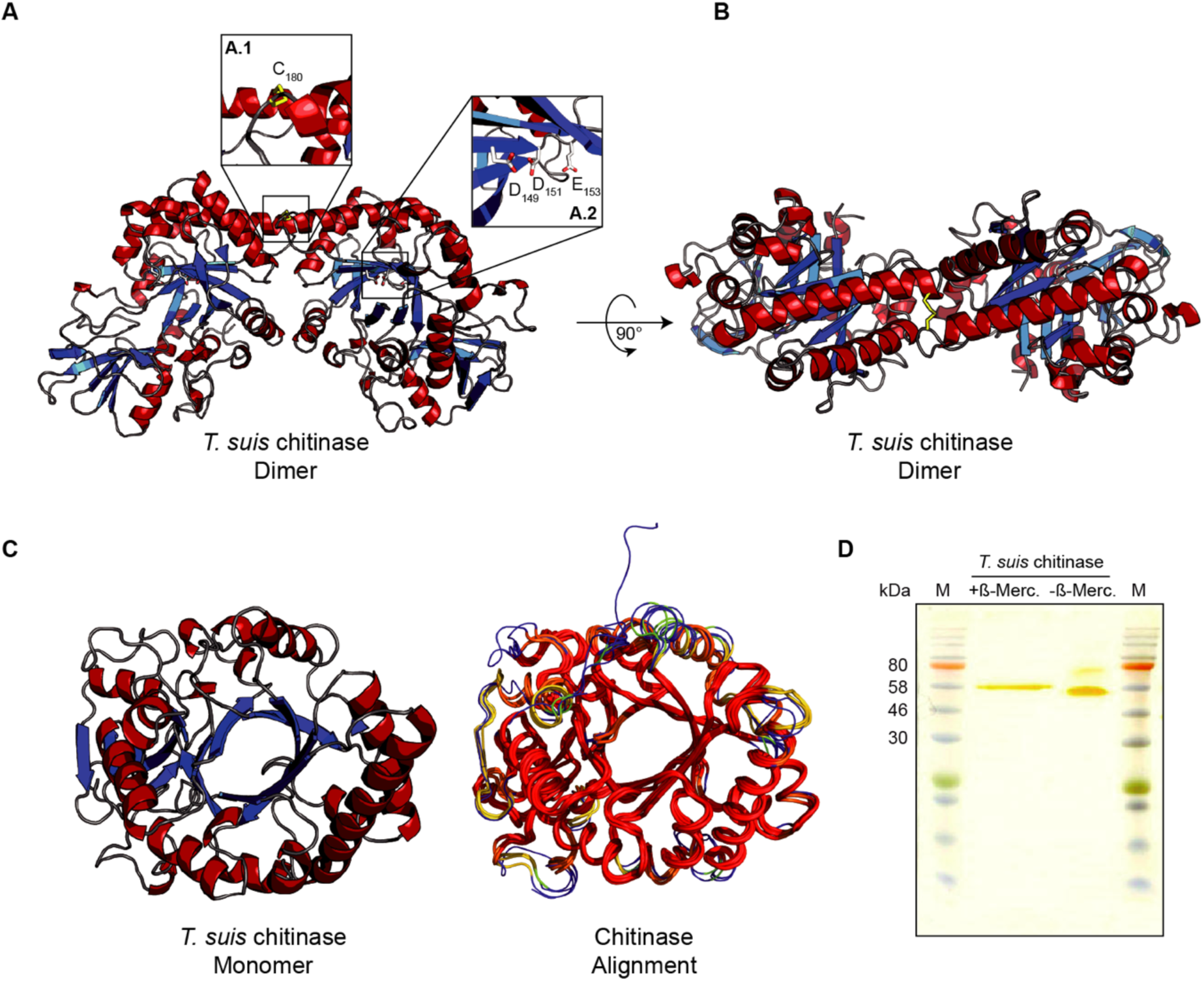
Three-dimensional structure of *T. suis* chitinase (*Ts*_Chit). (**A**) Overall structure of *Ts*-Chit forming a C2-symmetric dimer [α-helices in red, ß-strands in blue] The inset **A.1** depicts the dimer-stabilizing disulphide bond-forming cysteine residue at position 180, while the inset **A.2** highlights the catalytic DxDxE center. (**B**) *Ts*-Chit dimer rotated by 90° compared to (A). (**C**) *Ts*-Chit monomer and its structural alignment with AMCase (PDB Id 3FXY, (59)), human Chit1 (PDB-Id 1GUV, (60)), human YKL-40 (PDB-Id 1NWR, (61)) and murine Ym1 (PDB-Id 1VF8, (62)). (**D**) Silver stained SDS/PAGE gel of recombinantly expressed *Ts*-Chit (1 µg) under reducing (+ß-Merc.) and non-reducing (-ß-Merc.) loading conditions.

The structural und functional similarity of *T. suis* chitinase and murine AMCase could either mean that *Ts*-Chit is sensed similar to murine AMCase resulting in cumulative effects, or in turn, that *Ts*-Chit interferes with murine AMCase function. Following the idea that *T. suis* interferes with murine chitinase activity during allergic airway hyperreactivity, we more specifically compared surface features of *Ts*-Chit and mouse AMCase (PDB: 3FY1), the only acidic mouse chitinase with known experimental 3D structure. To that effect, the surface electrostatics for mammalian AMCase and the *Ts*-Chit monomer were visualized and appraised. While investigating the monomer alone did not reveal any strikingly identical surface patches, looking at the dimerization region around α3/α4-α3’/α4’ (α3:~127-142; α4:~161-180) revealed similarities with an acidic surface patch of AMCase (**Fig. 6A**).

**Figure 6.**
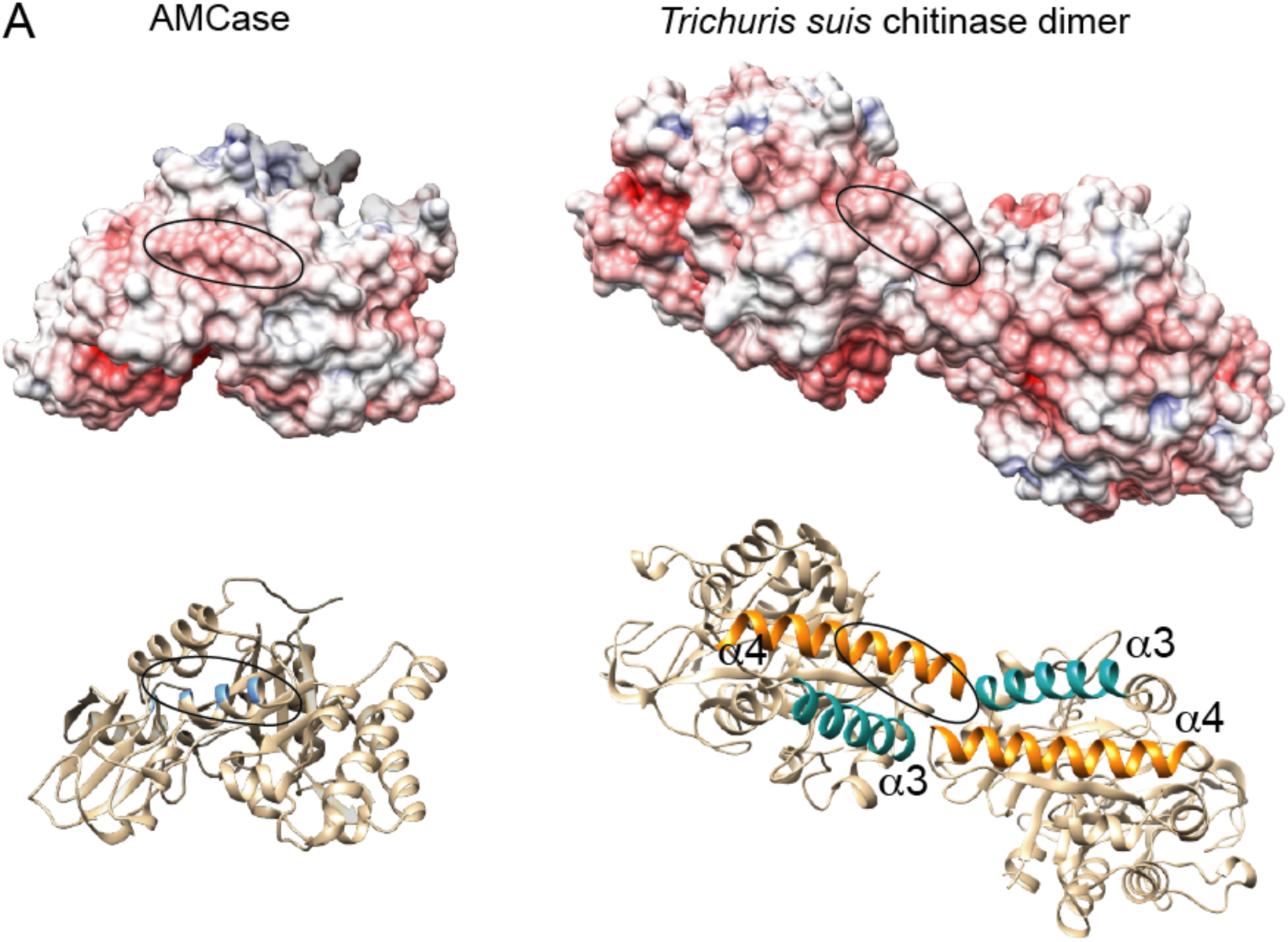
Electrostatic surface representation of murine AMCase (PDB 3FY1) and the *Ts*-Chit dimer. (**A**) Upper panel: In the *Ts*-Chit dimer, a surface patch of similar charge and shape as observed in murine AMCase is formed. The similar, acidic surface patch in both molecules is circled. Lower panel: Cartoon rendering of murine AMCase and *Ts*-Chit in the same orientation as in the upper panel. The location of the circled residues in AMCase are highlighted in blue. Helices α3 and α4 contributing to the dimer formation in *Ts*-Chit are colored turquoise and orange, respectively. The surface electrostatics were calculated with APBS and visualized with UCSF Chimera.

In summary, this first crystallographic structure analysis of a parasitic nematode chitinase revealed *Ts*-Chit dimerization and the concomitant existence of surface patches absent in the monomeric protein. The surface electrostatics of these dimer-specific patches show a remarkable similarity between *T. suis* chitinase and murine AMCase.

### *T. suis* chitinase interferes with murine chitinase in AHR

Considering the structural similarities, we asked whether *Ts*-Chit treatment of allergic mice interfered with the effector functions of its murine counterpart, AMCase. And indeed, chitinase activity was found to be reduced in the BAL fluid of allergic mice treated with intact and inactivated *Ts*-Chit compared to untreated (OVA) mice (**Fig. 7A**). Similarly, we detected decreased AMCase transcript levels (*Chia1*) mRNA transcripts in lung tissues of allergic *Ts*-Chit treated compared untreated (OVA) mice (**Fig. 7B**).

**Figure 7.**
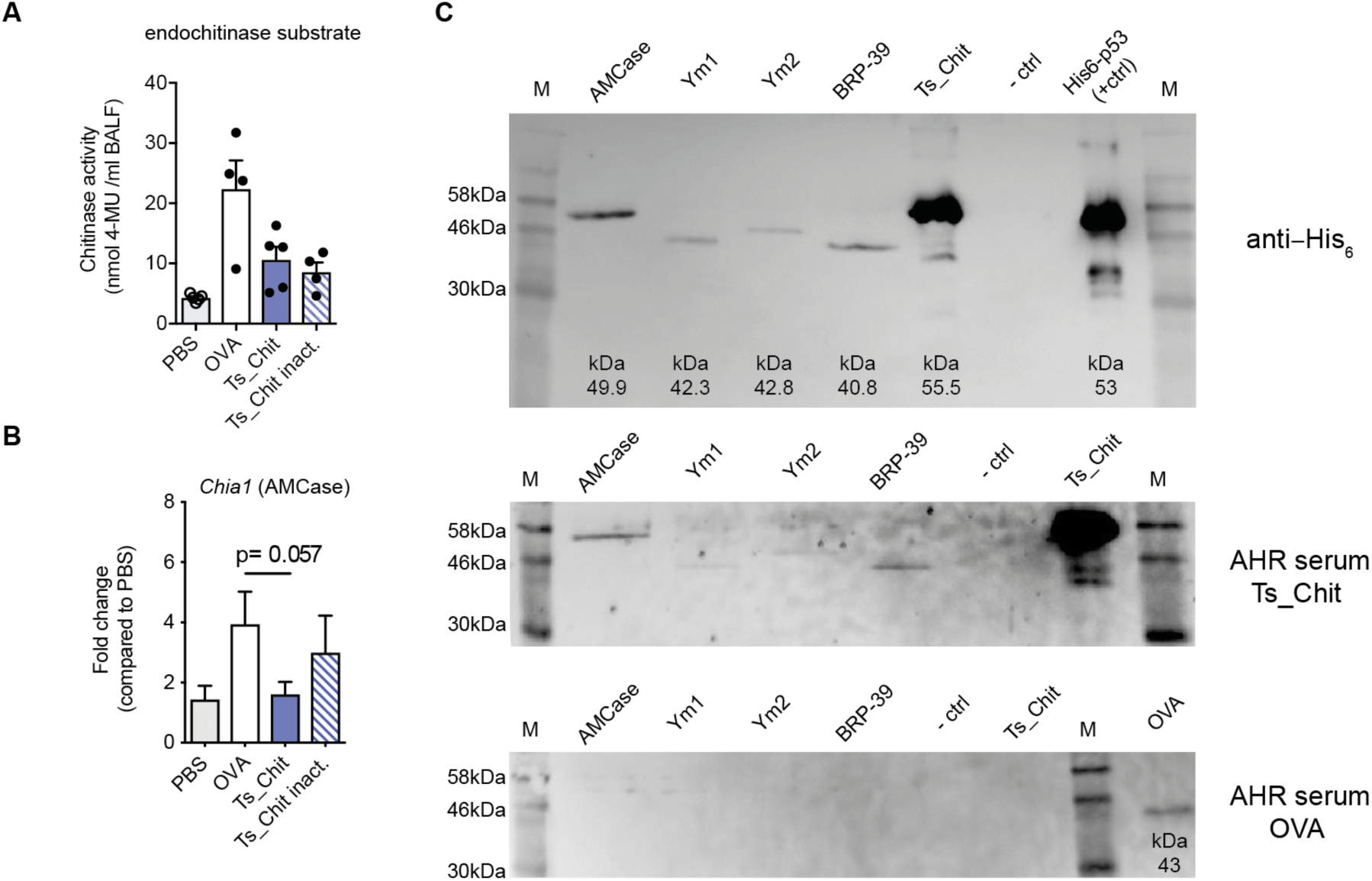
*Ts*-Chit treatment results in antibody responses against murine AMCase and BRP-39 and interferes with murine chitinase activity. (A) Chitinase assay testing BAL samples of allergic mice with endochitinase substrate. Data shown are mean + SEM of *n* = 4– 5 replicates. (**B**) mRNA expression of murine chitinase AMCase *(Chia1)* in lung tissue samples of mice from AHR challenge experiments after lung lavage. Expression was normalized by using the housekeeper gene *peptidyl prolylisomerase A* (*PPIA*) and is presented as fold difference to the PBS group. Data are presented as mean + SEM of *n* = 12 animals from *n* = 3 separate experiments. Data were analysed statistically using an unpaired *t*-test. (**C**) Western blot of recombinantly expressed murine AMCase, murine CLPs and *Ts*-Chit. Top blot of SDS/PAGE shows detection of recombinant proteins by anti-His_6_ antibody (1:200) after loading 20 µl of AMCase-, Ym1-, Ym2-, and BRP-39-transfected COS7 cell supernatants, 0.125 µg *Ts*-Chit and 0.2 µg p53 control protein. As negative control, supernatants of non-transfected COS7 cells were used (-ctrl). Middle blot shows murine chitinase and CLP loading, but incubation with serum antibodies from *Ts*-Chit treated, OVA-sensitized and challenged mice (1:100) and detection using anti-mouse IgG-HRP (1:2500). Bottom blot shows identical loading including one lane for OVA protein (0.5 µg) as positive control and incubation with serum antibodies from untreated, allergic (OVA) mice (1:100) detected using anti-mouse IgG-HRP (1:2500). For serum incubation, sera of *n* = 4 mice were pooled and representative Western blots from *n* = 3 separate experiments are shown.

Since chitinase inhibition by either administration of anti-AMCase sera (63), RNA interference (64) or small molecule inhibitors e.g. allosamidin (63,65) results in suppression of airway inflammation and hyperresponsiveness, we studied the serological response of cross-reactive anti-AMCase and anti-CLP antibodies. We speculated that the high degree of structural similarity between *T. suis* chitinase and murine chitinases or CLPs probably contributes to the development of cross-reactive antibodies when applied during allergic sensitization in murine AHR. Blotting recombinantly expressed murine AMCase and the CLPs Ym1, Ym2 and BRP-39 and probing the blots with AHR sera of *Ts*-Chit treated, allergic mice clearly demonstrated the presence of antibodies recognizing *Ts*-Chit but also murine AMCase and BRP-39 proteins (**Fig. 7C**). In turn, the serum of untreated, allergic mice recognized OVA protein only, but no chitinases or CLPs (**Fig. 7C**). Together, these data indicate a functional role for cross-reactive antibodies to murine AMCase and BRP-39 that are induced by *Ts*-Chit treatment.

## Discussion

Parasitic nematodes use a range of different mechanisms to not only invade their hosts, but also promote survival, growth and reproduction. A very elegant strategy involves immunomodulation of the host comprising of suppression, regulation or modulation of the host immune response. Being equipped with excretory and secretory systems (66), parasitic nematodes release numerous possible mediators into their environment that are most likely involved in communicating with the host and shaping its immune response (1). It is those yet unidentified ES immunomodulatory molecules that bear the therapeutic potential for targeting immune-mediated diseases.

The life cycle of many parasitic nematodes, however, is complex and involves uptake of infective larvae, hatching, host invasion or migration, growth and maturation, several molting events, mating and sexual reproduction. Accordingly, there is evidence for life cycle stage specific expression and release of distinct sets of ES molecules needed to fulfill those specific tasks (30,67,68). We have previously reported that total ES proteins of the very early larval stage (L1) of the whipworm *T. suis* very efficiently ameliorate allergic lung disease in a model of OVA-induced AHR when applied during the sensitization phase (33). Notably, in TSO therapy, patients are repeatedly exposed to the early larval stages of *T. suis* owing to the fact that *T. suis* only transiently infects human intestines (23,69).

This led us to suggest a role for mediators being released very early during infection by the first larval stages. In this study, we therefore focused on the identification of specific L1 ES immunomodulators of *T. suis* and their immunomodulatory mechanisms. We compared ES proteins collected from hatched L1 *versus* later larval stages (10, 18 and 28 day old) isolated from the intestines of infected pigs by using LC-MS/MS. Among the 33 proteins that were identified in the ES of *in vitro* hatched *T. suis* L1, 8 proteins were also found in the ES of L2 larvae, but no overlap to later larval stages. We concentrated on proteins selectively expressed by the first two larval stages (L1 and L2) and chose targets based on the presence of a signal peptide and the absence of a transmembrane domain. However, we recognize that a surprisingly large proportion of *T. suis* ES proteins are not encoded with a signal peptide, comparable with reports from other species (70–72). Over the course of this study, the genome of the porcine whipworm *T. suis* published in 2014 (73) had been complemented with two slightly larger assemblies of male and female worms, that in turn added on to the number of identified protein IDs when reanalyzing our MS data. Hence, ongoing data base curation of the *T. suis* genome can increase the number of potential immunomodulatory proteins in ES products.

Screening six recombinantly expressed L1 ES proteins, eukaryotically-produced in *L. tarentolae*, in a model of OVA-induced allergic airway hyperreactivity revealed that the application of *T. suis* KFD48490.1 markedly reduced inflammatory lung infiltrates, partly mimicking the effects of total *T. suis* L1 ES (33). Our results further proved that KFD48490.1 is an enzymatically active chitinase of *T. suis*. The enzyme retained its high activity in the pH range from 4 to 6, and thus is relatively resistant to acidic pH similar to acidic mammalian chitinases of humans and mice (74,75).

Chitinases are glycosyl hydrolases (GH) with the ability to directly degrade chitin polymers to low molecular weight chitooligomers (reviewed in (76)). Based on amino acid sequence similarity, the *T. suis* chitinase belongs to GH family 18 characterized by an enzyme core of 8 β-strands forming a barrel that is surrounded by α-helices, as well as the conserved sequence DxDxE forming the active site. Chitinases have essential roles in the life cycle of many parasites, supported by the existence of multiple genes as well as tissue and stage-specific expression patterns (77–79). Chitinase activity in helminth biology is thought to be essential during embryonic development, larval molting and degradation of the chitinous matrix (80,81), suggesting that chitinases might be attractive intervention targets. RNAi experiments demonstrated that inhibition of helminth chitinase prevented larval molting and hatching of microfilaria, but also induced the death of female worms of the rodent filarial nematode *Acanthocheilonema viteae* (49).

In the context of allergic lung disease, however, observing an active chitinase to mediate suppression of AHR was rather surprising. In recent years, studies in patients and animal models have shown that mammalian chitinases play a key role in mediating the Th2 driven inflammatory responses commonly associated with asthma and have been investigated to indicate disease severity (63,82,83). Mice express two enzymatically active chitinases, acidic mammalian chitinase (AMCase) and chitotriosidase (Chit) and a set of enzymatically inactive chitinase-like proteins (Ym1, Ym2, BRP39). AMCase is secreted by macrophages and epithelial cells of lung and gut. In the lungs, it is constitutively expressed and secreted into the airway lumen. STAT6-activating signals, such as IL-13, further induce AMCase expression and elevated AMCase mRNA and protein levels are detected in BAL of OVA-induced asthmatic mice (63). This latter study also showed that administration of anti-AMCase sera reduced BAL and tissue eosinophilia and BAL chitinase activity. Similarly, in a gut inflammation model it has been shown that the pan-chitinase inhibitor caffeine was able to dampen inflammation in a DSS-induced colitis model (84).

Here, we demonstrated that treating mice with a chitinase of *T. suis* (*Ts*-Chit) during sensitization reduced BAL eosinophilia, increased numbers of Relmα^+^ interstitial cells, improved lung function and reduced chitinase activity in BAL. Interestingly, heat-inactivated *Ts*-Chit was able to partly phenocopy these effects *in vivo*. We have no evidence for active *Ts*-Chit to induce Foxp3^+^ Treg cells, to interfere with antigen-specific and -unspecific T cell proliferation, to alter co-stimulatory molecule expression of antigen-presenting cells or induce NO in macrophages as shown in other studies (30,57). Yet, we cannot rule out any direct cellular effects of *Ts*-Chit on mammalian cells. Our *in vivo* results, however, suggest a mechanism related to the anti-AMCase-sera treatment of OVA-induced allergic mice reported by Zhu and colleagues (63). Similar to that study, we show here that *Ts*-Chit treatment had no suppressive effects on Th2 inductive responses such IL-4, IL-13 as well as IL-5 levels in BAL. Zhu and coworkers further demonstrated that anti-AMCase sera diminished the ability of IL-13 to stimulate eotaxin and monocyte-chemotactic protein 1 (MCP-1), an effect that was induced by the worm chitinase in the present study. Besides anti-AMCase treatment, other approaches have been used to address AMCase activity, such as inhibition with small molecule (65) or RNA interference (64), and demonstrated suppressed eotaxin, MCP-1 and MIP-1ß levels when AMCase activity or expression was inhibited.

The underlying mechanisms on the ability of AMCase to promote chemokine and cytokine expression have been investigated by Hartl and coworkers (85) who showed that AMCase physically interacts with epidermal growth factor receptor (EGFR) to induce the production of e.g. MCP-1, CCL17, and IL-8 in lung epithelial cells. The authors also reported an autocrine and/or paracrine feedback mechanism for AMCase in protecting airway epithelial cells from Fas ligand (FasL)- and growth factor withdrawal-induced apoptosis (85,86).

Sequence alignment demonstrates a relatively high degree of similarity between *T. suis* chitinase and AMCase, but also to CLPs such as YM1, Ym2 or BRP39. Given this homology of *T. suis* and murine chitinases and our *in vivo* observations on reduced chemokine expression and eosinophil infiltration, but non-suppressed Th2 responses, it is tempting to speculate that *Ts*-Chit administration during the sensitization phase induces cross-reactive antibodies that inhibit murine chitinases and/or CLPs. *Ts*-Chit treatment clearly induced an antibody response against murine AMCase and BRP-39 in our model of OVA-induced allergic asthma; additionally, we observed decreased chitinase activity in BAL and lung tissue decrease in AMCase transcripts. Whether decreased chitinase activity and reduced AMCase transcription are a consequence of the overall reduced lung inflammation or result from chitinase protein interference still need to be verified.

The X-ray crystal structure of *Ts*-Chit reported here confirmed the expected high similarity of the nematode chitinase with murine acidic mammalian chitinase. In addition, analysis of surface electrostatics of the *Ts*-Chit dimer suggests the existence of an acidic surface patch on the nematode protein that bears similarity to a surface feature on AMCase monomers. This suggests that the response elicited by *Ts*-Chit may be due to molecular mimicry; however, future studies need to establish whether this is indeed the case. Site-directed mutagenesis within the dimerization region or the acidic surface patch will be needed to address the impact those structural features to the formation of cross-reactive antibodies.

Molecular mimicry is a phenomenon that is triggered by a high degree of similarity of pathogen and host epitopes and has previously been described for helminth defense molecules of *Fasciola hepatica* that mimic the molecular function of host antimicrobial molecules (87) or the TGF-ß mimic secreted by *Heligmosomoides polygyrus* with profound Treg-inducing capacities (19). To the best of our knowledge, a pathogen-driven molecular mimicry inducing antibodies that interfere with Th2-dependent host chitinase activity has not been described so far.

An interesting thought that arises when considering the induction of cross-reactive antibodies under inflammatory conditions, is the heterologous response of patients receiving TSO therapy in published reports of multiple sclerosis (MS) or UC trials (28,88). Considering that a certain threshold of inflammation is required to trigger polyclonal B cell responses and induction of cross-reactive antibodies, it is conceivable that, depending on the individual inflammation score and its kinetics, TSO induces a more or less anti-inflammatory treatment response.

In summary, our study investigated L1 stage-specific, secreted proteins of *T. suis* and their capacity for immunomodulation in treating allergic lung disease. We identified an active chitinase of *T. suis* that, when applied during sensitization of an allergic response, reduced eosinophilia and lung chemokine responses. These findings add to our understanding of specific functions of early secreted *T. suis* products in helminth biology and when considered as therapeutics to create an anti-inflammatory environment. The structural similarity between the worm chitinase and its host equivalent, the absence of direct cellular effects and the presence of cross-reactive antibodies potentially interfering with murine chitinase activity suggests an alternative mechanism of *Ts*-Chit ameliorating OVA-induced, allergic lung disease that likely involves structural mimicry. Such a mechanism of immunomodulation could be one of those unrecognized strategies of mediating anti-inflammatory responses that explain the degree of variation among the responses of individual TSO-treated subjects in clinical trials.

## Acknowledgements

The authors gratefully thank Marion Müller, Christiane Palissa, Yvonne Weber, Bettina Sonnenburg und Beate Anders for excellent technical assistance and Franziska Rohr for her help within the animal facility. We also thank Prof. Judith E. Allen for her expertise, critical advice and intellectual discussion of the topic. This work was supported by the German Research Foundation (GRK 1673 to S. H.), the FAZIT-STIFTUNG (to K. B.) and Coronado Biosciences (to S. H.).

## Author contribution (CRediT taxonomy)

Conceptualization, F. E. and S. H.; Methodology, K. B., F. E., M. S. W., P. H. M., J. Z., K. J.; Investigation, K. B., F. E., A. A. K., K. J., A.N., M. S. W., P. H. M., A. H.; Data Curation, K. J., A. N., P. H. M.; Writing, F. E., K. B. and S. H.; Funding Acquisition, S. H.; Resources, T. E. S., A. H., M. S. W., J. Z., S. H.; Formal Analysis, A. H., A. A. K., A. H.; Visualization, F. E., K. B., P. H. M., M. S. W., A. H., Supervision, F. E., M. S. W. and S. H.

## Nonstandard abbreviations

AAM: alternatively activated macrophages
AHR: airway hyperreactivity
AMCase: acidic mammalian chitinase
BAL: bronchoalveolar lavage
Chit: Chitinase
CLPs: Chitinase-like proteins
ES: excretory-secretory
ESP: excretory-secretory proteins
GH: glycosyl hydrolases
L1: fist larval stage
LEXSY: Leishmania expression system
MS: mass spectrometry
OVA: Ovalbumin
SPs: soluble proteins
TCEP: Tris(2-carboxyethyl)phosphine
*T. suis*: *Trichuris suis*
*Ts*-Chit: *Trichuris suis* chitinase
*Ts*-ES: *Trichuris suis* excretory-secretory proteins
TSO: *Trichuris suis* ova
TTO: *Trichuris trichiura* ova
UC: ulcerative colitis

---

**Table S1:**
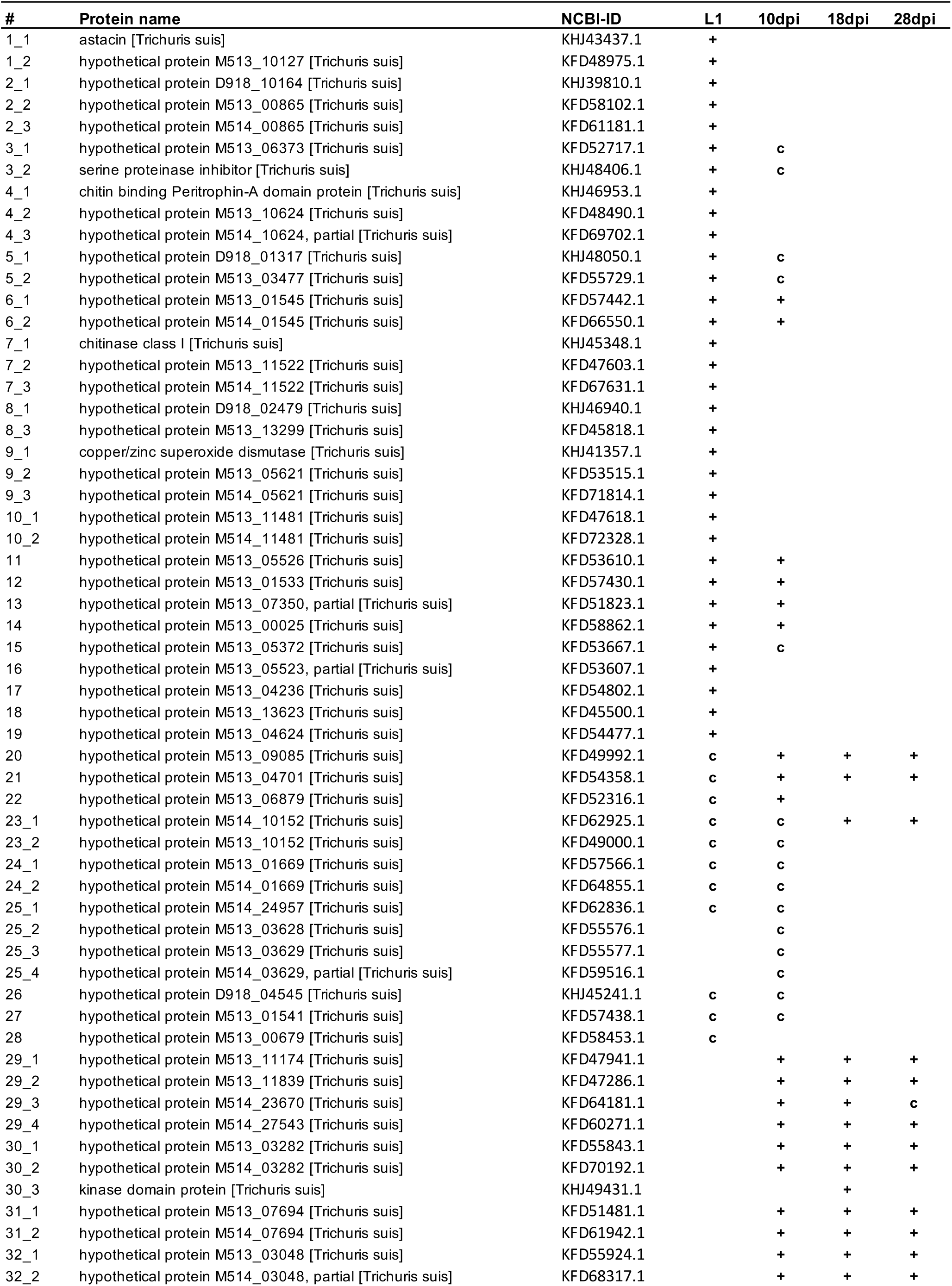
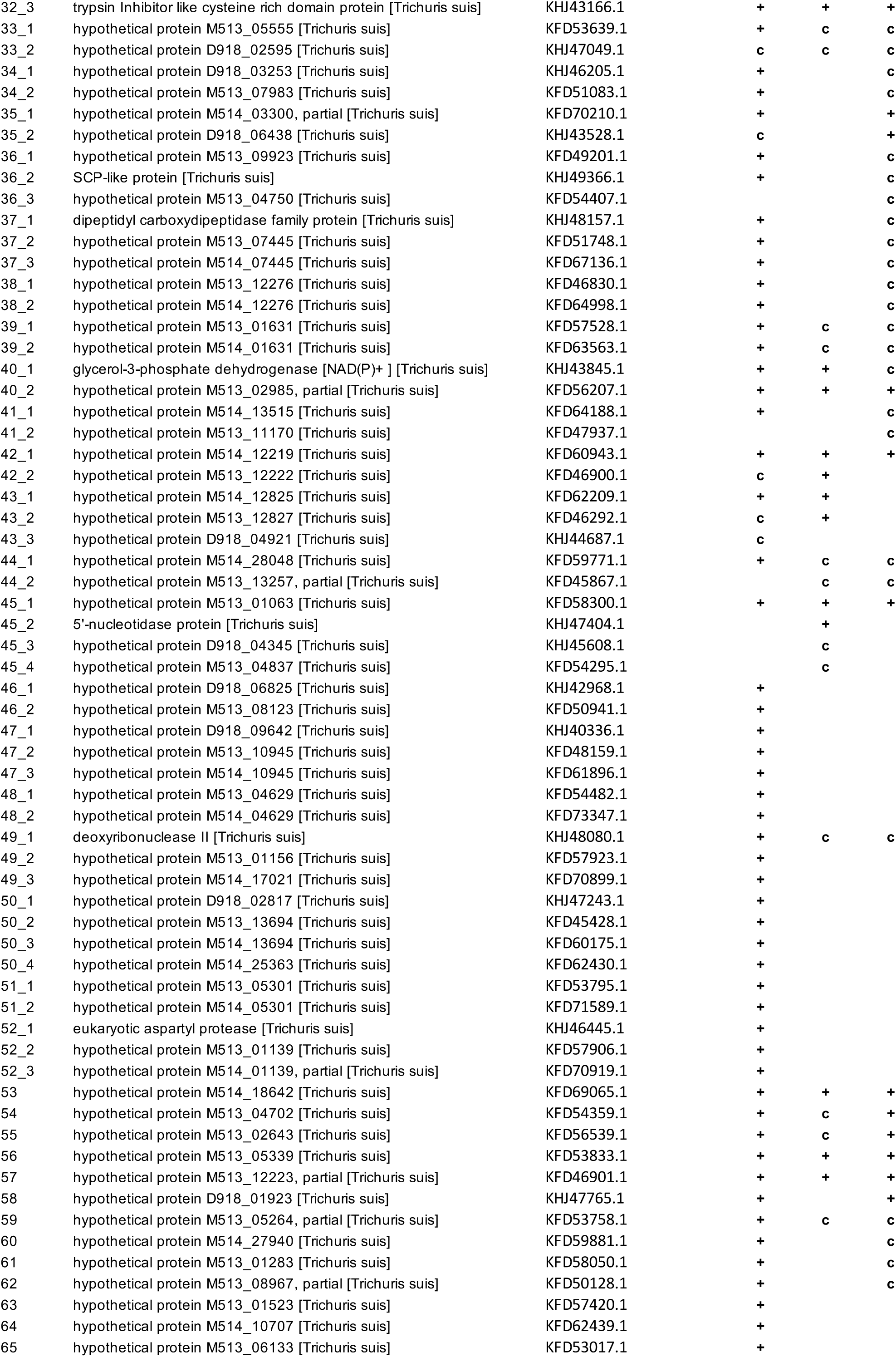
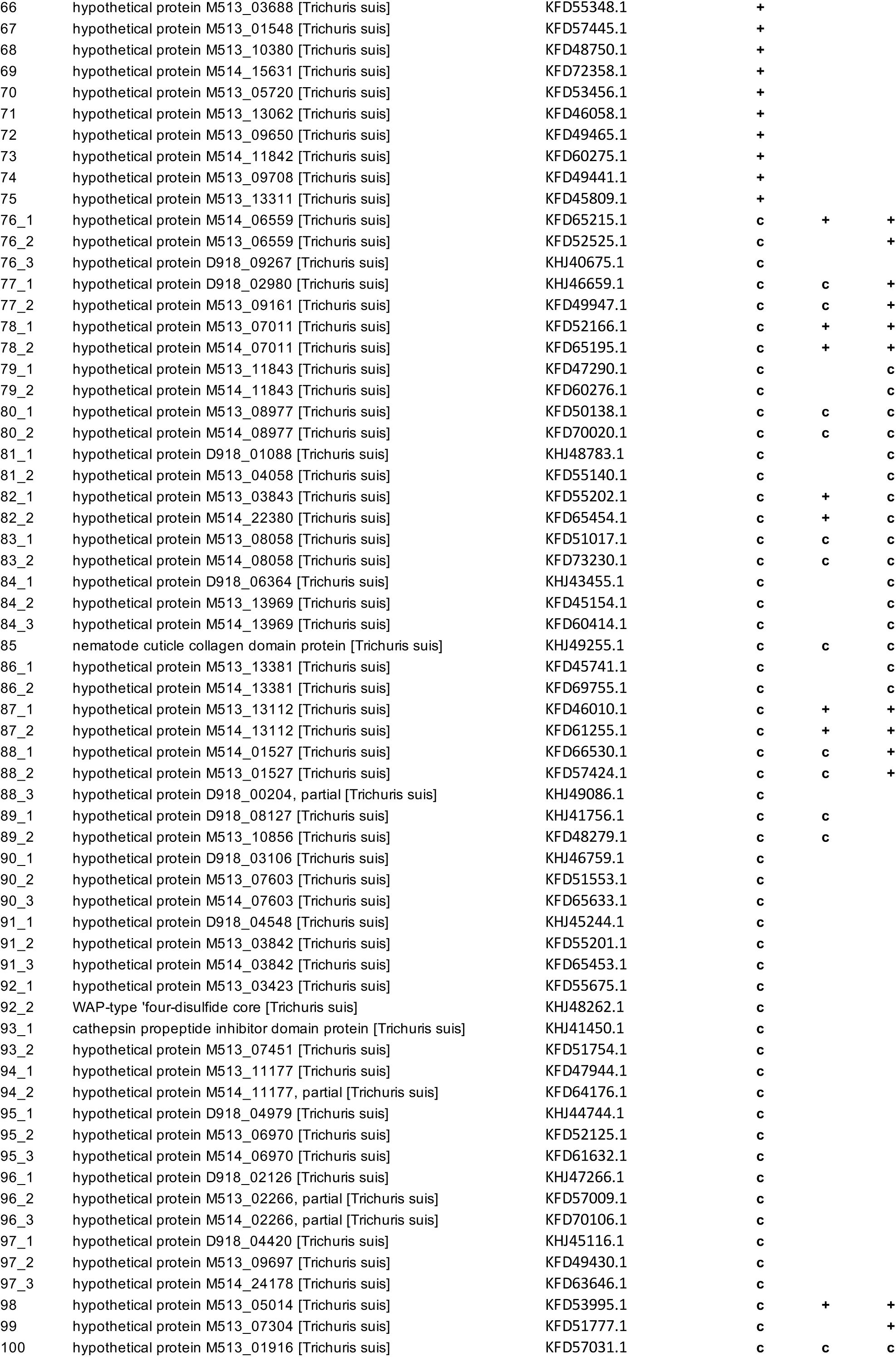
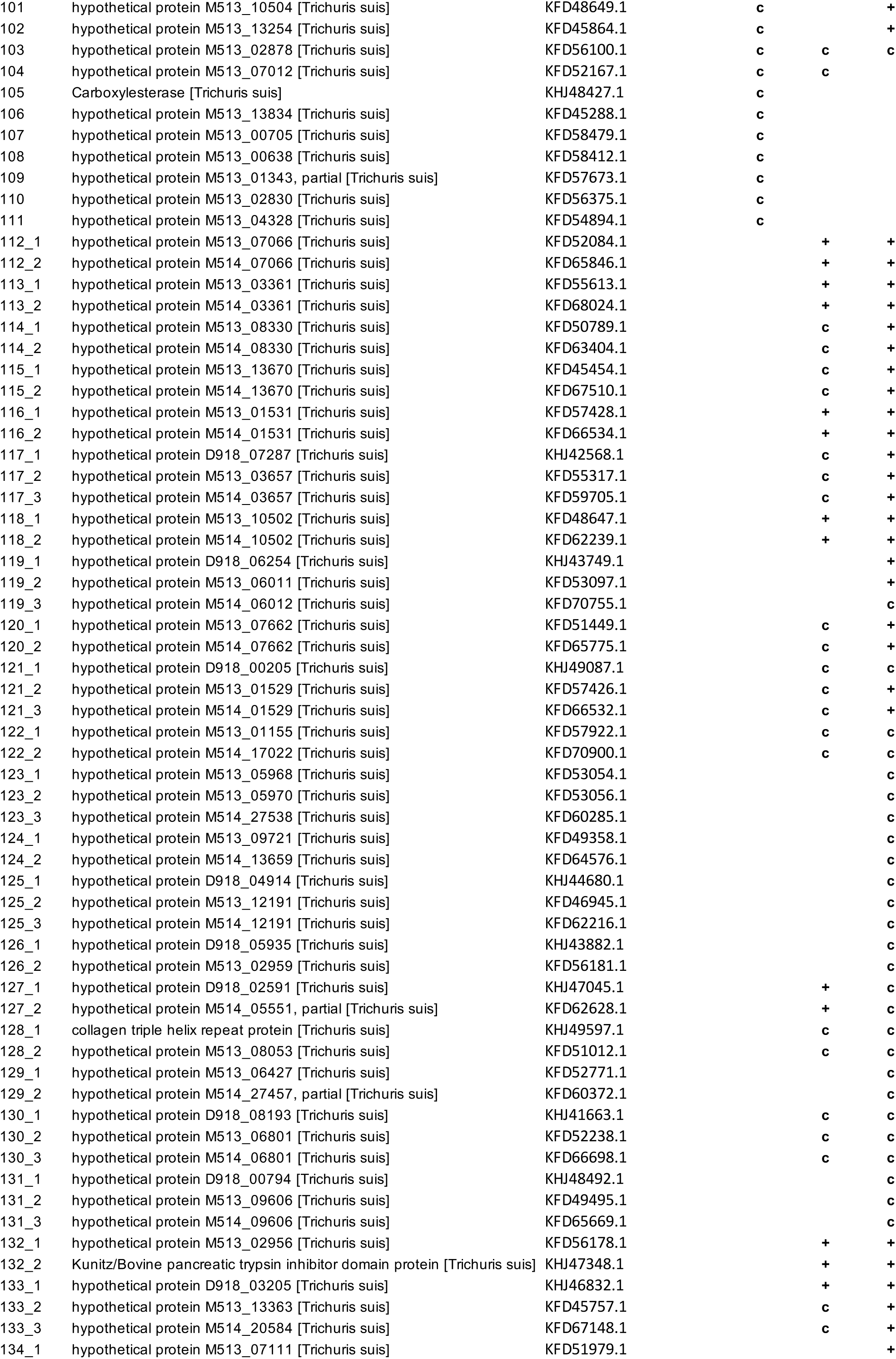
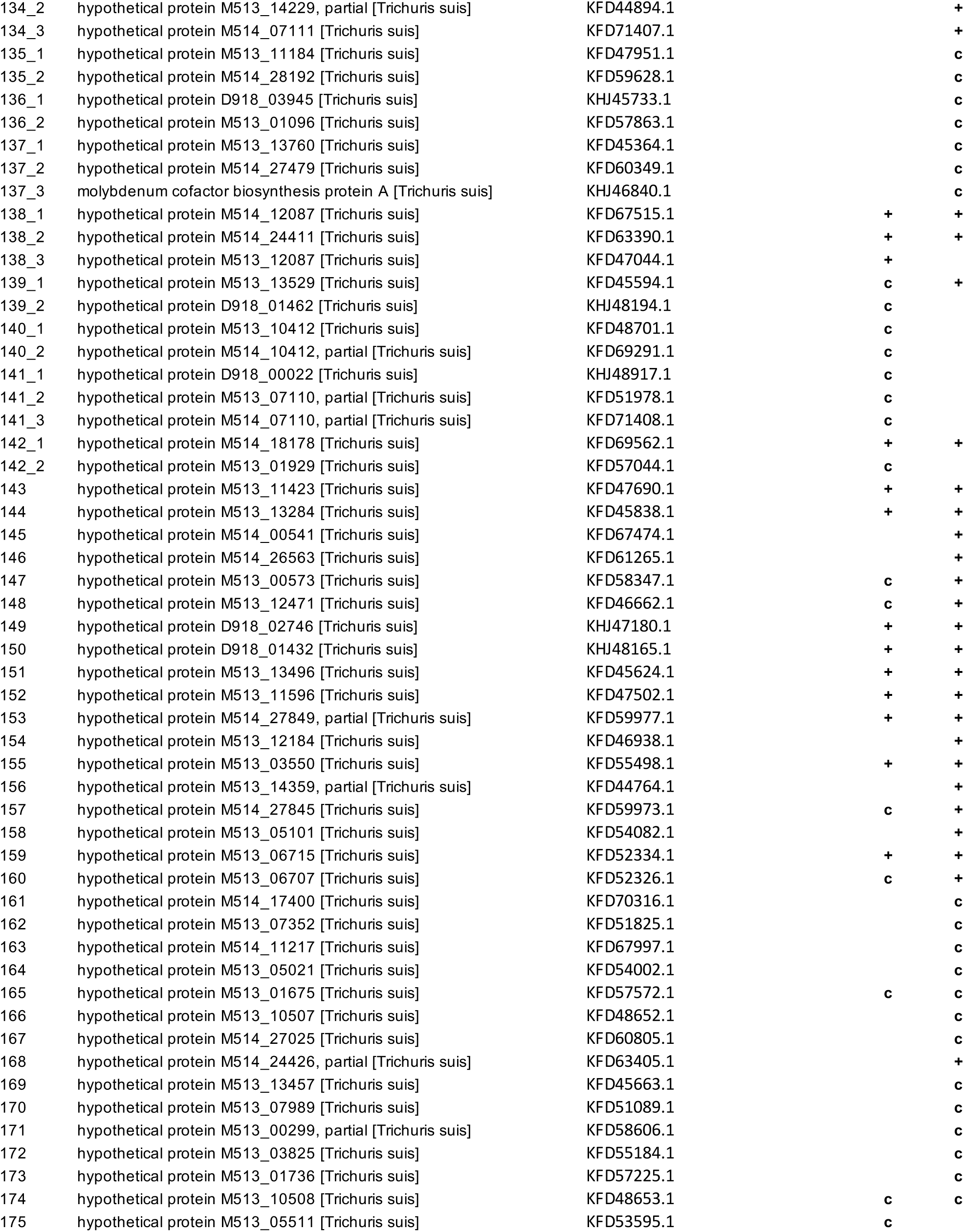
Mass spectrometric identification of *T. suis* larvae ES proteins. Precipitated proteins were reduced, alkylated, tryptic digested and analyzed by LC-MSMS and Mascot searches against NCBI database. Significant hits (+) were identified with ≥ 2 peptides (p< 0.01), non-significant candidates (c) are also listed. Homologous proteins are grouped to the same #-number.

## Supplementary Information S2

**Figure S2.**
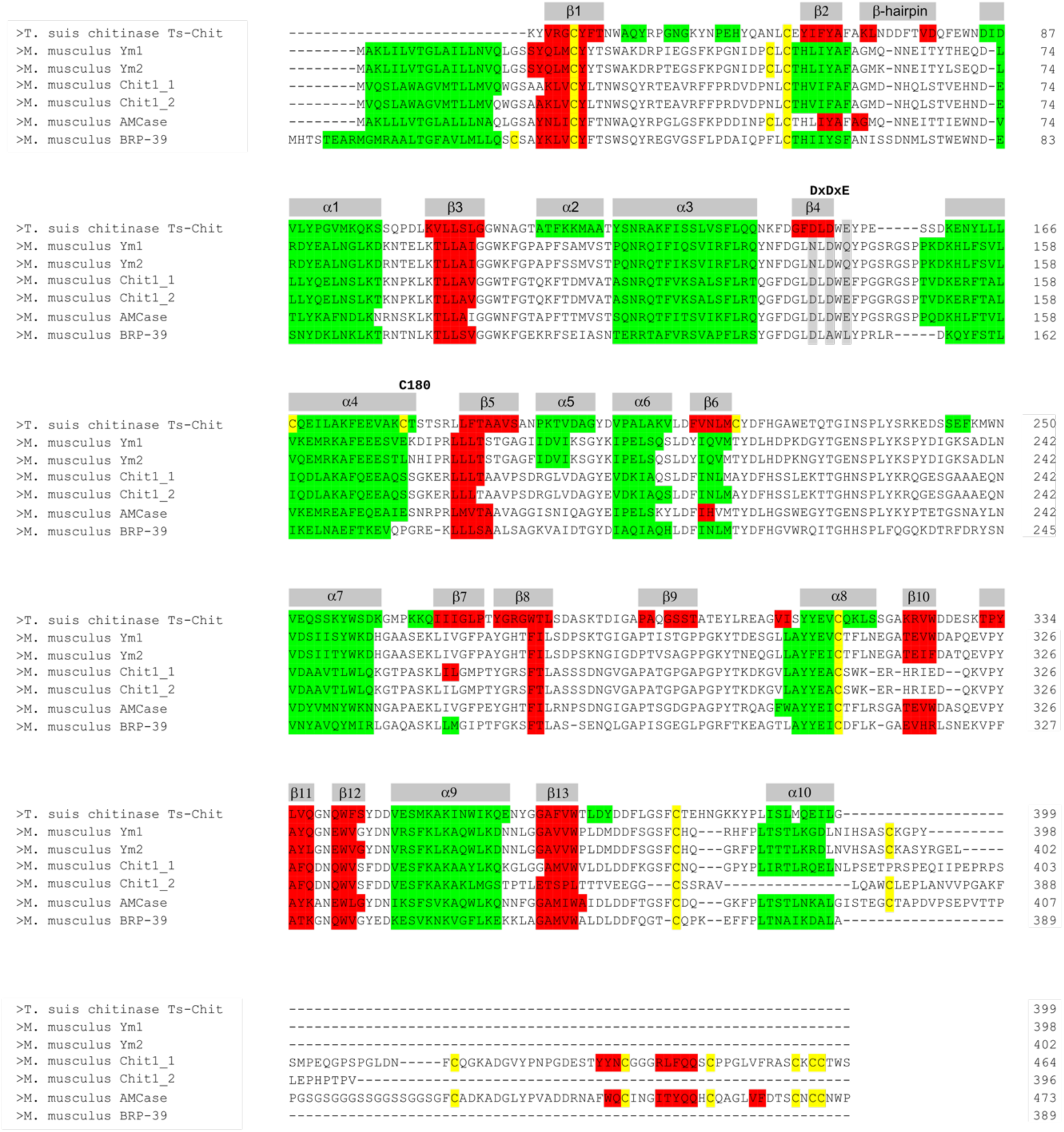
Comparison of topology and secondary structure elements of *T. suis* chitinase with mouse chitinases and chitinase like proteins (CLPs). Structure-based amino acid sequence alignment of *T. suis* chitinase (*Ts*-Chit) with murine Ym1 (*Chil3*), Ym2 (*Chil4*), chitotriosidase isoforms 1 & 2 (*Chit1_1, Chit1_2*), AMCase (*Chia*) and BRP-39 (*Chil1*). Secondary structure elements were predicted and visualized with PSIPRED [1] and SBAL [2], indicating α-helices in green, β-strands in red and cysteines in yellow. The top line shows the annotation with secondary structure elements observed in the crystal structure of *Ts*-Chit and the catalytic motive DxDxE.

## References

1. Hewitson JP, Grainger JR, Maizels RM. Helminth immunoregulation: The role of parasite secreted proteins in modulating host immunity. Mol Biochem Parasitol. 2009 Sep;167(1–9):1–11.

2. McSorley HJ, Hewitson JP, Maizels RM. Immunomodulation by helminth parasites: Defining mechanisms and mediators. International Journal for Parasitology. 2013 Mar 1;43(3):301–10.

3. Coakley G, Buck AH, Maizels RM. Host parasite communications—Messages from helminths for the immune system. Mol Biochem Parasitol. 2016 Jul;208(1):33–40.

4. Schnoeller C, Rausch S, Pillai S, Avagyan A, Wittig BM, Loddenkemper C, et al. A Helminth Immunomodulator Reduces Allergic and Inflammatory Responses by Induction of IL-10-Producing Macrophages. J Immunol. 2008 Mar 15;180(6):4265–72.

5. Daniłowicz-Luebert E, Steinfelder S, Kühl AA, Drozdenko G, Lucius R, Worm M, et al. A nematode immunomodulator suppresses grass pollen-specific allergic responses by controlling excessive Th2 inflammation. Int J Parasitol. 2013 Mar;43(3–4):201–10.

6. Whelan RA, Rausch S, Ebner F, Günzel D, Richter JF, Hering NA, et al. A Transgenic Probiotic Secreting a Parasite Immunomodulator for Site-Directed Treatment of Gut Inflammation. Molecular Therapy. 2014 Jul 2;

7. Ziegler T, Rausch S, Steinfelder S, Klotz C, Hepworth MR, Kühl AA, et al. A novel regulatory macrophage induced by a helminth molecule instructs IL-10 in CD4+ T cells and protects against mucosal inflammation. J Immunol. 2015 Feb 15;194(4):1555–64.

8. Pineda MA, Lumb F, Harnett MM, Harnett W. ES-62, a therapeutic anti-inflammatory agent evolved by the filarial nematode Acanthocheilonema viteae. Mol Biochem Parasitol. 2014 Apr;194(1–2):1–8.

9. Rodgers DT, McGrath MA, Pineda MA, Al-Riyami L, Rzepecka J, Lumb F, et al. The Parasitic Worm Product ES-62 Targets Myeloid Differentiation Factor 88–Dependent Effector Mechanisms to Suppress Antinuclear Antibody Production and Proteinuria in MRL/lpr Mice. Arthritis Rheumatol. 2015 Apr;67(4):1023–35.

10. McInnes IB, Leung BP, Harnett M, Gracie JA, Liew FY, Harnett W. A Novel Therapeutic Approach Targeting Articular Inflammation Using the Filarial Nematode-Derived Phosphorylcholine-Containing Glycoprotein ES-62. The Journal of Immunology. 2003 Aug 15;171(4):2127–33.

11. Rzepecka J, Siebeke I, Coltherd JC, Kean DE, Steiger CN, Al-Riyami L, et al. The helminth product, ES-62, protects against airway inflammation by resetting the Th cell phenotype. Int J Parasitol. 2013 Mar;43(3–4):211–23.

12. Zaccone P, Fehérvári Z, Jones FM, Sidobre S, Kronenberg M, Dunne DW, et al. Schistosoma mansoni antigens modulate the activity of the innate immune response and prevent onset of type 1 diabetes. Eur J Immunol. 2003 May;33(5):1439–49.

13. Zaccone P, Burton O, Miller N, Jones FM, Dunne DW, Cooke A. Schistosoma mansoni egg antigens induce Treg that participate in diabetes prevention in NOD mice. European Journal of Immunology. 2009 Apr 1;39(4):1098–107.

14. Janssen L, Silva Santos GL, Muller HS, Vieira ARA, de Campos TA, de Paulo Martins V. Schistosome-Derived Molecules as Modulating Actors of the Immune System and Promising Candidates to Treat Autoimmune and Inflammatory Diseases. J Immunol Res. 2016;2016.

15. Sarazin A, Dendooven A, Delbeke M, Gatault S, Pagny A, Standaert A, et al. Treatment with P28GST, a schistosome-derived enzyme, after acute colitis induction in mice: Decrease of intestinal inflammation associated with a down regulation of Th1/Th17 responses. PLoS One. 2018 Dec 28;13(12).

16. Ramos-Benitez MJ, Ruiz-Jimenez C, Rosado-Franco JJ, Ramos-Pérez WD, Mendez LB, Osuna A, et al. Fh15 Blocks the Lipopolysaccharide-Induced Cytokine Storm While Modulating Peritoneal Macrophage Migration and CD38 Expression within Spleen Macrophages in a Mouse Model of Septic Shock. mSphere. 2018 Dec 19;3(6).

17. Roig J, Saiz ML, Galiano A, Trelis M, Cantalapiedra F, Monteagudo C, et al. Extracellular Vesicles From the Helminth Fasciola hepatica Prevent DSS-Induced Acute Ulcerative Colitis in a T-Lymphocyte Independent Mode. Front Microbiol. 2018 May 23;9. 18.

18. McSorley HJ, Blair NF, Smith KA, McKenzie ANJ, Maizels RM. Blockade of IL-33 release and suppression of type 2 innate lymphoid cell responses by helminth secreted products in airway allergy. Mucosal Immunol. 2014 Feb 5;

19. Johnston CJC, Smyth DJ, Kodali RB, White MPJ, Harcus Y, Filbey KJ, et al. A structurally distinct TGF-β mimic from an intestinal helminth parasite potently induces regulatory T cells. Nat Commun. 2017 Nov 23;8.

20. Togre NS, Bhoj PS, Khatri VK, Tarnekar A, Goswami K, Shende MR, et al. SXP-RAL Family Filarial Protein, rWbL2, Prevents Development of DSS-Induced Acute Ulcerative Colitis. Indian J Clin Biochem. 2018 Jul;33(3):282–9.

21. Amdare NP, Khatri VK, Yadav RSP, Tarnekar A, Goswami K, Reddy MVR. Therapeutic potential of the immunomodulatory proteins Wuchereria bancrofti L2 and Brugia malayi abundant larval transcript 2 against streptozotocin-induced type 1 diabetes in mice. Journal of Helminthology. 2017 Sep;91(5):539–48.

22. Fleming JO, Weinstock JV. Clinical trials of helminth therapy in autoimmune diseases: rationale and findings. Parasite Immunology. 2015;37(6):277–92.

23. Summers RW, Elliott DE, Qadir K, Urban JF Jr, Thompson R, Weinstock JV. Trichuris suis seems to be safe and possibly effective in the treatment of inflammatory bowel disease. Am J Gastroenterol. 2003 Sep;98(9):2034–41.

24. Summers RW, Elliott DE, Urban JF, Thompson RA, Weinstock JV. Trichuris suis therapy for active ulcerative colitis: A randomized controlled trial. Gastroenterology. 2005 Apr 1;128(4):825–32.

25. Summers RW, Elliott DE, Urban JF, Thompson R, Weinstock JV. Trichuris suis therapy in Crohn’s disease. Gut. 2005 Jan 1;54(1):87–90.

26. Schölmerich J, Fellermann K, Seibold FW, Rogler G, Langhorst J, Howaldt S, et al. A Randomised, Double-blind, Placebo-controlled Trial of Trichuris suis ova in Active Crohn’s Disease. J Crohns Colitis. 2017 Apr;11(4):390–9.

27. Hollander E, Uzunova G, Taylor BP, Noone R, Racine E, Doernberg E, et al. Randomized crossover feasibility trial of helminthic Trichuris suis ova versus placebo for repetitive behaviors in adult autism spectrum disorder. The World Journal of Biological Psychiatry. 2018 Sep 19;0(0):1–9.

28. Fleming J, Hernandez G, Hartman L, Maksimovic J, Nace S, Lawler B, et al. Safety and efficacy of helminth treatment in relapsing-remitting multiple sclerosis: Results of the HINT 2 clinical trial. Mult Scler. 2019 Jan 1;25(1):81–91.

29. Beer RJS. Studies on the biology of the life-cycle of Trichuris suis Schrank, 1788. Parasitology. 1973 Dec;67(3):253–62.

30. Leroux L-P, Nasr M, Valanparambil R, Tam M, Rosa BA, Siciliani E, et al. Analysis of the Trichuris suis excretory/secretory proteins as a function of life cycle stage and their immunomodulatory properties. Scientific Reports. 2018 Oct 29;8(1):15921.

31. Kuijk LM, Klaver EJ, Kooij G, van der Pol SMA, Heijnen P, Bruijns SCM, et al. Soluble helminth products suppress clinical signs in murine experimental autoimmune encephalomyelitis and differentially modulate human dendritic cell activation. Molecular Immunology. 2012 Jun;51(2):210–8.

32. Hansen TVA, Williams AR, Denwood M, Nejsum P, Thamsborg SM, Friis C. Pathway of oxfendazole from the host into the worm: Trichuris suis in pigs. International Journal for Parasitology: Drugs and Drug Resistance. 2017 Dec 1;7(3):416–24.

33. Ebner F, Hepworth MR, Rausch S, Janek K, Niewienda A, Kühl A, et al. Therapeutic potential of larval excretory/secretory proteins of the pig whipworm Trichuris suis in allergic disease. Allergy. 2014 Jul 29;

34. Klaver EJ, Kuijk LM, Laan LC, Kringel H, van Vliet SJ, Bouma G, et al. Trichuris suis-induced modulation of human dendritic cell function is glycan-mediated. International Journal for Parasitology. 2013 Mar;43(3–4):191–200.

35. Ottow MK, Klaver EJ, van der Pouw Kraan TCTM, Heijnen PD, Laan LC, Kringel H, et al. The helminth Trichuris suis suppresses TLR4-induced inflammatory responses in human macrophages. Genes Immun. 2014 Oct;15(7):477–86.

36. Hoeksema MA, Laan LC, Postma JJ, Cummings RD, de Winther MPJ, Dijkstra CD, et al. Treatment with Trichuris suis soluble products during monocyte-to-macrophage differentiation reduces inflammatory responses through epigenetic remodeling. FASEB J. 2016;30(8):2826–36.

37. Laan LC, Williams AR, Stavenhagen K, Giera M, Kooij G, Vlasakov I, et al. The whipworm (Trichuris suis) secretes prostaglandin E2 to suppress proinflammatory properties in human dendritic cells. FASEB J. 2017 Feb;31(2):719–31.

38. Mueller U, Förster R, Hellmig M, Huschmann FU, Kastner A, Malecki P, et al. The macromolecular crystallography beamlines at BESSY II of the Helmholtz-Zentrum Berlin: Current status and perspectives. Eur Phys J Plus. 2015 Jul 22;130(7):141.

39. Sparta KM, Krug M, Heinemann U, Mueller U, Weiss MS. XDSAPP2.0. J Appl Cryst. 2016 Jun 1;49(3):1085–92.

40. Fadel F, Zhao Y, Cousido-Siah A, Ruiz FX, Mitschler A, Podjarny A. X-Ray Crystal Structure of the Full Length Human Chitotriosidase (CHIT1) Reveals Features of Its Chitin Binding Domain. PLoS One. 2016 Apr 25;11(4).

41. Emsley P, Lohkamp B, Scott WG, Cowtan K. Features and development of Coot. Acta Crystallogr D Biol Crystallogr. 2010 Apr 1;66(Pt 4):486–501.

42. Murshudov GN, Vagin AA, Dodson EJ. Refinement of Macromolecular Structures by the Maximum-Likelihood Method. Acta Cryst D. 1997 May 1;53(3):240–55.

43. Bryson K, McGuffin LJ, Marsden RL, Ward JJ, Sodhi JS, Jones DT. Protein structure prediction servers at University College London. Nucleic Acids Res. 2005 Jul 1;33(Web Server issue):W36–8.

44. Wang CK, Broder U, Weeratunga SK, Gasser RB, Loukas A, Hofmann A. SBAL: a practical tool to generate and edit structure-based amino acid sequence alignments. Bioinformatics. 2012 Apr 1;28(7):1026–7.

45. Lobley A, Sadowski MI, Jones DT. pGenTHREADER and pDomTHREADER: new methods for improved protein fold recognition and superfamily discrimination. Bioinformatics. 2009 Jul 15;25(14):1761–7.

46. Jurrus E, Engel D, Star K, Monson K, Brandi J, Felberg LE, et al. Improvements to the APBS biomolecular solvation software suite. Protein Sci. 2018;27(1):112–28.

47. Pettersen EF, Goddard TD, Huang CC, Couch GS, Greenblatt DM, Meng EC, et al. UCSF Chimera--a visualization system for exploratory research and analysis. J Comput Chem. 2004 Oct;25(13):1605–12.

48. Dahiya N, Tewari R, Hoondal GS. Biotechnological aspects of chitinolytic enzymes: a review. Appl Microbiol Biotechnol. 2006 Aug 1;71(6):773–82.

49. Tachu B, Pillai S, Lucius R, Pogonka T. Essential Role of Chitinase in the Development of the Filarial Nematode Acanthocheilonema viteae. Infect Immun. 2008 Jan;76(1):221–8.

50. Gooyit M, Tricoche N, Lustigman S, Janda KD. Dual Protonophore–Chitinase Inhibitors Dramatically Affect O. volvulus Molting. J Med Chem. 2014 Jul 10;57(13):5792–9.

51. Sanders NL, Mishra A. Role of Interleukin-18 in the Pathophysiology of Allergic Diseases. Cytokine Growth Factor Rev. 2016 Dec;32:31–9.

52. Nair MG, Du Y, Perrigoue JG, Zaph C, Taylor JJ, Goldschmidt M, et al. Alternatively activated macrophage-derived RELM-α is a negative regulator of type 2 inflammation in the lung. The Journal of Experimental Medicine. 2009 Apr 13;206(4):937.

53. Sutherland TE, Rückerl D, Logan N, Duncan S, Wynn TA, Allen JE. Ym1 induces RELMα and rescues IL-4Rα deficiency in lung repair during nematode infection. PLOS Pathogens. 2018 Nov 30;14(11):e1007423.

54. Wolfs IMJ, Stöger JL, Goossens P, Pöttgens C, Gijbels MJJ, Wijnands E, et al. Reprogramming macrophages to an anti-inflammatory phenotype by helminth antigens reduces murine atherosclerosis. The FASEB Journal. 2013 Sep 16;28(1):288–99.

55. Paterson JCM, Garside P, Kennedy MW, Lawrence CE. Modulation of a Heterologous Immune Response by the Products of Ascaris suum. Infect Immun. 2002 Nov;70(11):6058–67.

56. Hartmann S, Kyewski B, Sonnenburg B, Lucius R. A filarial cysteine protease inhibitor down-regulates T cell proliferation and enhances interleukin-10 production. Eur J Immunol. 1997 Sep;27(9):2253–60.

57. Grainger JR, Smith KA, Hewitson JP, McSorley HJ, Harcus Y, Filbey KJ, et al. Helminth secretions induce de novo T cell Foxp3 expression and regulatory function through the TGF-β pathway. J Exp Med. 2010 Oct 25;207(11):2331–41.

58. Sutherland TE, Maizels RM, Allen JE. Chitinases and chitinase-like proteins: potential therapeutic targets for the treatment of T-helper type 2 allergies. Clinical & Experimental Allergy. 2009;39(7):943–55.

59. Olland AM, Strand J, Presman E, Czerwinski R, Joseph-McCarthy D, Krykbaev R, et al. Triad of polar residues implicated in pH specificity of acidic mammalian chitinase. Protein Sci. 2009 Mar;18(3):569–78.

60. Fusetti F, Moeller H von, Houston D, Rozeboom HJ, Dijkstra BW, Boot RG, et al. Structure of Human Chitotriosidase IMPLICATIONS FOR SPECIFIC INHIBITOR DESIGN AND FUNCTION OF MAMMALIAN CHITINASE-LIKE LECTINS. J Biol Chem. 2002 Jul 12;277(28):25537–44.

61. Fusetti F, Pijning T, Kalk KH, Bos E, Dijkstra BW. Crystal Structure and Carbohydrate-binding Properties of the Human Cartilage Glycoprotein-39. J Biol Chem. 2003 Sep 26;278(39):37753–60.

62. Tsai M-L, Liaw S-H, Chang N-C. The crystal structure of Ym1 at 1.31Å resolution. Journal of Structural Biology. 2004 Dec 1;148(3):290–6.

63. Zhu Z, Zheng T, Homer RJ, Kim Y-K, Chen NY, Cohn L, et al. Acidic Mammalian Chitinase in Asthmatic Th2 Inflammation and IL-13 Pathway Activation. Science. 2004 Jun 11;304(5677):1678–82.

64. Yang C-J, Liu Y-K, Liu C-L, Shen C-N, Kuo M-L, Su C-C, et al. Inhibition of acidic mammalian chitinase by RNA interference suppresses ovalbumin-sensitized allergic asthma. Hum Gene Ther. 2009 Dec;20(12):1597–606.

65. Matsumoto T, Inoue H, Sato Y, Kita Y, Nakano T, Noda N, et al. Demethylallosamidin, a chitinase inhibitor, suppresses airway inflammation and hyperresponsiveness. Biochemical and Biophysical Research Communications. 2009 Dec 4;390(1):103–8.

66. Basyoni MMA, Rizk EMA. Nematodes ultrastructure: complex systems and processes. J Parasit Dis. 2016 Dec;40(4):1130–40.

67. Wang T, Van Steendam K, Dhaenens M, Vlaminck J, Deforce D, Jex AR, et al. Proteomic analysis of the excretory-secretory products from larval stages of Ascaris suum reveals high abundance of glycosyl hydrolases. PLoS Negl Trop Dis. 2013;7(10):e2467.

68. Hewitson JP, Harcus Y, Murray J, van Agtmaal M, Filbey KJ, Grainger JR, et al. Proteomic analysis of secretory products from the model gastrointestinal nematode Heligmosomoides polygyrus reveals dominance of Venom Allergen-Like (VAL) proteins. J Proteomics. 2011 Aug 24;74(9):1573–94.

69. Kradin RL, Badizadegan K, Auluck P, Korzenik J, Lauwers GY. Iatrogenic Trichuris suis Infection in a Patient With Crohn Disease. Archives of Pathology & Laboratory Medicine Online. 2009.

70. Chehayeb JF, Robertson AP, Martin RJ, Geary TG. Proteomic analysis of adult Ascaris suum fluid compartments and secretory products. PLoS Negl Trop Dis. 2014 Jun;8(6):e2939.

71. Hewitson JP, Harcus YM, Curwen RS, Dowle AA, Atmadja AK, Ashton PD, et al. The secretome of the filarial parasite, Brugia malayi: Proteomic profile of adult excretory–secretory products. Molecular and Biochemical Parasitology. 2008 Jul 1;160(1):8–21.

72. Cass CL, Johnson JR, Califf LL, Xu T, Hernandez HJ, Stadecker MJ, et al. Proteomic Analysis of Schistosoma mansoni Egg Secretions. Mol Biochem Parasitol. 2007 Oct;155(2):84–93.

73. Jex AR, Nejsum P, Schwarz EM, Hu L, Young ND, Hall RS, et al. Genome and transcriptome of the porcine whipworm Trichuris suis. Nat Genet. 2014 Jul;46(7):701–6.

74. Boot RG, Blommaart EFC, Swart E, Vlugt KG der, Bijl N, Moe C, et al. Identification of a Novel Acidic Mammalian Chitinase Distinct from Chitotriosidase. J Biol Chem. 2001 Mar 2;276(9):6770–8.

75. Okawa K, Ohno M, Kashimura A, Kimura M, Kobayashi Y, Sakaguchi M, et al. Loss and Gain of Human Acidic Mammalian Chitinase Activity by Nonsynonymous SNPs. Mol Biol Evol. 2016 Dec;33(12):3183–93.

76. Hamid R, Khan MA, Ahmad M, Ahmad MM, Abdin MZ, Musarrat J, et al. Chitinases: An update. J Pharm Bioallied Sci. 2013;5(1):21–9.

77. Wu Y, Egerton G, Underwood AP, Sakuda S, Bianco AE. Expression and Secretion of a Larval-specific Chitinase (Family 18 Glycosyl Hydrolase) by the Infective Stages of the Parasitic Nematode, Onchocerca volvulus. J Biol Chem. 2001 Nov 9;276(45):42557–64.

78. Wu Y, Adam R, Williams SA, Bianco AE. Chitinase genes expressed by infective larvae of the filarial nematodes, Acanthocheilonema viteae and Onchocerca volvulus. Molecular and Biochemical Parasitology. 1996 Jan 1;75(2):207–19.

79. Fuhrman JA, Lane WS, Smith RF, Piessens WF, Perler FB. Transmission-blocking antibodies recognize microfilarial chitinase in brugian lymphatic filariasis. PNAS. 1992 Mar 1;89(5):1548–52.

80. Adam R, Kaltmann B, Rudin W, Friedrich T, Marti T, Lucius R. Identification of Chitinase as the Immunodominant Filarial Antigen Recognized by Sera of Vaccinated Rodents. J Biol Chem. 1996 Jan 19;271(3):1441–7.

81. Ward KA, Fairbairn D. Chitinase in Developing Eggs of Ascaris suum (Nematoda). The Journal of Parasitology. 1972;58(3):546–9.

82. Zimmermann N, Mishra A, King NE, Fulkerson PC, Doepker MP, Nikolaidis NM, et al. Transcript Signatures in Experimental Asthma: Identification of STAT6-Dependent and - Independent Pathways. The Journal of Immunology. 2004 Feb 1;172(3):1815–24.

83. Zhao J, Zhu H, Wong CH, Leung KY, Wong WSF. Increased lungkine and chitinase levels in allergic airway inflammation: A proteomics approach. PROTEOMICS. 2005;5(11):2799–807.

84. Lee I-A, Low D, Kamba A, Llado V, Mizoguchi E. Oral caffeine administration ameliorates acute colitis by suppressing chitinase 3-like 1 expression in intestinal epithelial cells. J Gastroenterol. 2014 Aug;49(8):1206–16.

85. Hartl D, He CH, Koller B, Da Silva CA, Homer R, Lee CG, et al. Acidic Mammalian Chitinase Is Secreted via an ADAM17/Epidermal Growth Factor Receptor-dependent Pathway and Stimulates Chemokine Production by Pulmonary Epithelial Cells. J Biol Chem. 2008 Nov 28;283(48):33472–82.

86. Hartl D, He CH, Koller B, Da Silva CA, Kobayashi Y, Lee CG, et al. Acidic Mammalian Chitinase Regulates Epithelial Cell via a Chitinolytic-Independent Mechanism. J Immunol. 2009 Apr 15;182(8):5098–106.

87. Robinson MW, Donnelly S, Hutchinson AT, To J, Taylor NL, Norton RS, et al. A Family of Helminth Molecules that Modulate Innate Cell Responses via Molecular Mimicry of Host Antimicrobial Peptides. PLoS Pathog. 2011 May 12;7(5).

88. Huang X, Zeng L-R, Chen F-S, Zhu J-P, Zhu M-H. Trichuris suis ova therapy in inflammatory bowel disease. Medicine (Baltimore). 2018 Aug 24;97(34).

## References

1. Bryson K, McGuffin LJ, Marsden RL, Ward JJ, Sodhi JS, Jones DT. Protein structure prediction servers at University College London. Nucleic Acids Res. 2005;33: W36–W38.

2. Wang CK, Broder U, Weeratunga SK, Gasser RB, Loukas A, Hofmann A. SBAL: a practical tool to generate and edit structure-based amino acid sequence alignments. Bioinformatics. 2012;28: 1026–1027. doi:10.1093/bioinformatics/bts035

